# Gene regulation network inference using k-nearest neighbor-based mutual information estimation-Revisiting an old DREAM

**DOI:** 10.1101/2021.12.20.473242

**Authors:** Lior I. Shachaf, Elijah Roberts, Patrick Cahan, Jie Xiao

**Affiliations:** Department of Biophysics, Johns Hopkins University, 3400 N. Charles Street, Baltimore, MD, 21218, USA; 10x Genomics, 6230 Stoneridge Mall Road, Pleasanton, CA 94588-3260, USA; Institute for Cell Engineering, Department of Biomedical Engineering, Department of Molecular Biology and Genetics, Johns Hopkins School of Medicine, 733 N. Broadway, Baltimore, MD 21205, U. S. A.; Department of Biophysics and Biophysical Chemistry, Johns Hopkins School of Medicine, 725 N. Wolfe Street, WBSB 708, Baltimore, MD, 21205

**Keywords:** Gene regulatory network inference, mutual information, k-nearest neighbor

## Abstract

**Background:** A cell exhibits a variety of responses to internal and external cues. These responses are possible, in part, due to the presence of an elaborate gene regulatory network (GRN) in every single cell. In the past twenty years, many groups worked on reconstructing the topological structure of GRNs from large-scale gene expression data using a variety of inference algorithms. Insights gained about participating players in GRNs may ultimately lead to therapeutic benefits. Mutual information (MI) is a widely used metric within this inference/reconstruction pipeline as it can detect any correlation (linear and non-linear) between any number of variables (*n*-dimensions). However, the use of MI with continuous data (for example, normalized fluorescence intensity measurement of gene expression levels) is sensitive to data size, correlation strength and underlying distributions, and often requires laborious and, at times, *ad hoc* optimization.

**Results:** In this work, we first show that estimating MI of a bi- and tri-variate Gaussian distribution using *k*-nearest neighbor (kNN) MI estimation results in significant error reduction as compared to commonly used methods based on fixed binning. Second, we demonstrate that implementing the MI-based kNN Kraskov-Stoögbauer-Grassberger (KSG) algorithm leads to a significant improvement in GRN reconstruction for popular inference algorithms, such as Context Likelihood of Relatedness (CLR). Finally, through extensive *in-silico* benchmarking we show that a new inference algorithm CMIA (Conditional Mutual Information Augmentation), inspired by CLR, in combination with the KSG-MI estimator, outperforms commonly used methods.

**Conclusions:** Using three canonical datasets containing 15 synthetic networks, the newly developed method for GRN reconstruction - which combines CMIA, and the KSG-MI estimator - achieves an improvement of 20-35% in precision-recall measures over the current gold standard in the field. This new method will enable researchers to discover new gene interactions or choose gene candidates for experimental validations.

## Background

Most cells in a multicellular organism contain the same genome, yet they can differentiate into different cell types and adapt to different environmental conditions [1]. These responses to internal and external cues are possible due to the presence of an elaborate gene regulatory network (GRN). A GRN is the genome’s “flowchart” for various biological processes such as sensing, development, and metabolism, enabling the cell to follow specific instructions upon an internal or external stimulation. Understanding how genomic flowcharts are organized brings the potential to remediate dysfunctional ones [2] and design new ones for synthetic biology [3].

Advances in large-scale gene expression data collected from omic-level microarrays and RNA-seq experiments allow the construction of basic networks by clustering co-expressed genes using statistical correlation metrics such as covariance and threshold to determine the statistical significance [4]. Another common practice is to monitor the expression of multiple genes in response to perturbations and then infer the relationship between these genes [5]. Currently, there are several classes of methods to infer GRNs from expression data, such as the Bayesian networks method, the statistical/information theory method, and ordinary differential equations (ODEs) (see excellent reviews [6–8]).

Originally introduced for communication systems by Shannon in the late 40s [9], mutual information (MI) was quickly adopted by other disciplines as a statistical tool to evaluate the dependence between variables. Unlike the abovementioned traditional correlation methods like covariance, MI can detect linear and non-linear relationship between variables and can be applied to test the dependence between any number of variables (n-dimensions).

Over the last twenty years, researchers have implemented many methods employing MI to reconstruct GRNs, such as Relevance Networks [10]; ARACNE (Algorithm for the Reconstruction of Accurate Cellular Networks, [11]); and CLR (Context Likelihood of Relatedness, [12]). Using MI with two variables (*i.e.* genes) is straightforward, but due to the positive and symmetric nature of two-way MI [13], MI with only two variables cannot distinguish between direct and indirect regulation, coregulation, or logical gate-type interactions [14, 15]. To overcome these issues, a few groups have used different three-dimensional MI measures in inference algorithms [14, 16, 17] (for a comprehensive list of methods, see Mousavian *et al.* [18]). Importantly, in most methods using MI, continuous input (i.e., normalized fluorescence intensity data for gene expression) needs to be discretized first to build probability density functions (PDF). This practice is known to be sensitive to data size, correlation strength and underlying distributions [19].

In general, the simplest and most computationally inexpensive method to discretize continuous data is fixed (width) binning (FB) (**Fig. 1A**}, where a histogram with a fixed number of bins (or bin width) determined by certain statistical rules is used to model the PDF. For finite data size, FB generally under- or over-estimates MI (**Fig. S1A**). Over the years, researchers developed different methods to mitigate bin number sensitivity and to better estimate (or correct the bias in) MI, especially for data of small sizes. These methods correct either the entropies {Miller-Madow [20]) or the probability distribution by adaptive partitioning (AP) [21], k-Nearest Neighbor (kNN) [22] (**Fig. 1B**), kernel density estimator (KDE) [14] and/or B-spline functions, in which data points are divided into fractions between a predefined number of adjacent bins [23]. Unfortunately, all these methods make assumptions on the density distribution and require adjustment of parameters by the user for different scenarios except for kNN, which is shown to be accurate and robust across different values of *k* [19, 22]. However, kNN is rarely used due to the higher computational costs it entailed [24] or the limited improvement for two variables (2d) in downstream analysis.

**Figure 1.**
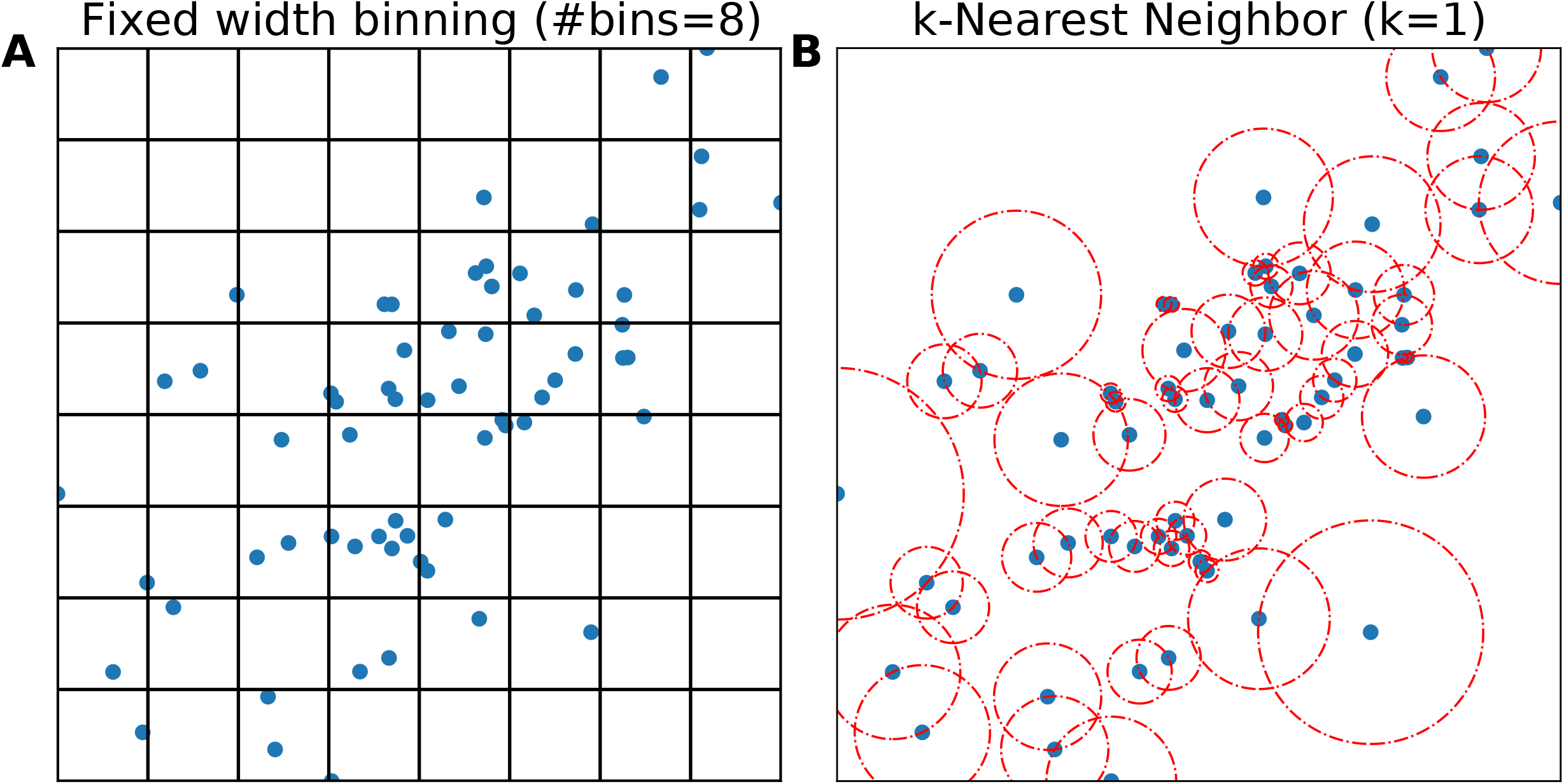
Illustration of two methods to evaluate distribution: (A) Fixed width binning, and (B) k-Nearest-Neighbor (k=1). Data points are shown as blue circles, bin edges are shown in black, and distances to k=1 neighbor as the radius of dashed red circles.

The problem of accurately estimating the correlation between genes has only worsened in this new era of single cell transcriptome studies, as data is larger yet sparser, often with non-Gaussian distributions. In this work, we focus on two subjects: (a) Improving MI estimation – we present an implementation of a three-way MI estimator based on kNN, which addresses large errors in estimating MI measures for three variables (3d). (b) Improving GRN inference – we present CMIA (Conditional Mutual Information Augmentation), a novel inference algorithm inspired by Synergy-Augmented CLR (SA-CLR) [17]. By testing various mutual information estimators against the ground truth solved from an analytical solution and comparing their performance using *in silico* GRN benchmarking data, we find that kNN-based three-way MI estimator Kraskov-Stoögbauer-Grassberger (KSG) improves the performance of common GRN inference methods. Together with the inference algorithm, CMIA, it outperforms other commonly used GRN reconstruction methods in the field.

## Results

### Benchmark MI estimations of a Gaussian distribution

To evaluate the performance of different mutual information (MI) estimators on continuous data, we calculated their deviations from the true underlying value by defining a percent error:

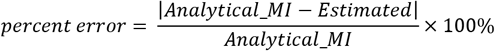

In most biologically relevant cases, one does not know what the true MI value is, because one does not know the probability distributions of the variables we are concerned with. Nevertheless, the true underlying value of MI of a few distributions such as Gaussian distribution can be analytically calculated. Therefore, to allow quantitative comparisons between different MI estimators, we used the analytical solution of Shannon’s entropy for a Gaussian distribution (see Additional file 1) to calculate the MI by entropy summation (**Table 1**). We then compared all methods of different data sizes (100, 1K, 10K, referring to the number of different conditions/perturbations/time points of individual genes) and different correlation strengths (0.3, 0.6, 0.9) between two or three variables (number of genes, 2d or 3d) drawn from a Gaussian distribution with a mean at zero and a variance of one (the absolute values of mean and variance are not important in the calculation as the final solution only contains correlation, see Additional file 1). For two-way MI (two variables, or 2d), we compared the following MI estimators: (i) Maximum Likelihood (ML, given by Shannon, Table 1), (ii) Miller-Madow correction (MM, see Additional file 1), (iii) Kozachenko-Leonenko (KL) [25], and (iv) KSG **(Fig. 2A**). The first two methods use FB to discretize the continuous data, and in general the best number of bins changes depending on the data size and correlation between variables (Additional file 2**: Fig. S1A**). As *a priori* the correlation strength is unknown, for the number of bins we used the common practice 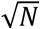, where N equals the number of data points, and the result was rounded down to align with methods in the next section. The latter two methods both use kNN, and we found that any selection of *k* resulted in good alignment with the analytical solution (see Additional file 2: **Fig. S1B**). We chose the third nearest-neighbor (*k*=3) as recommended by Kraskov et al [22] because a *k* value of 3 resulted in a good trade-off between precision and computational cost. As shown in **Fig. 2**, in all cases the two kNN-based MI estimators performed well similarly and outperformed the fixed-binning methods judged by the percentage error.

**Figure 2.**
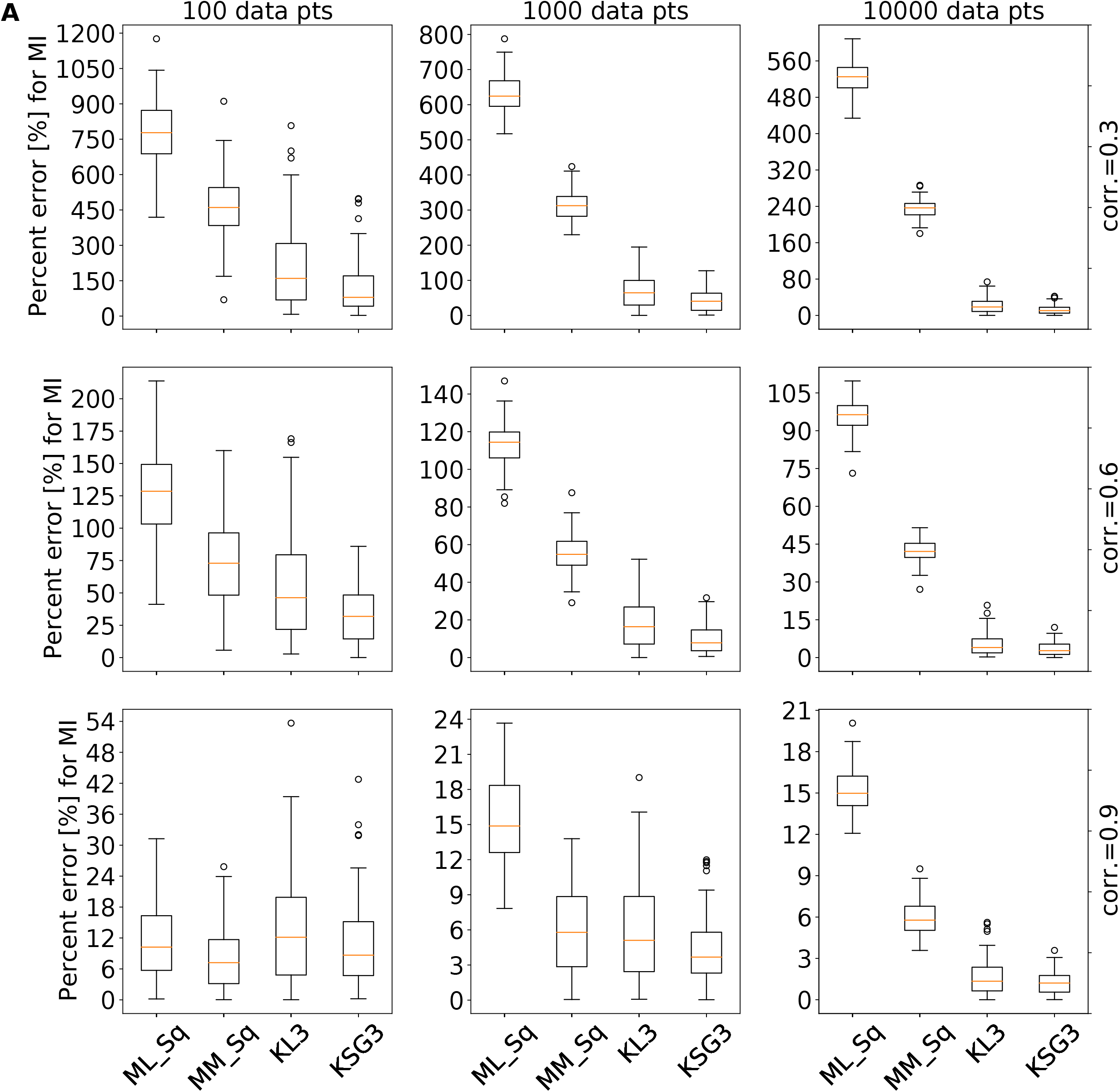
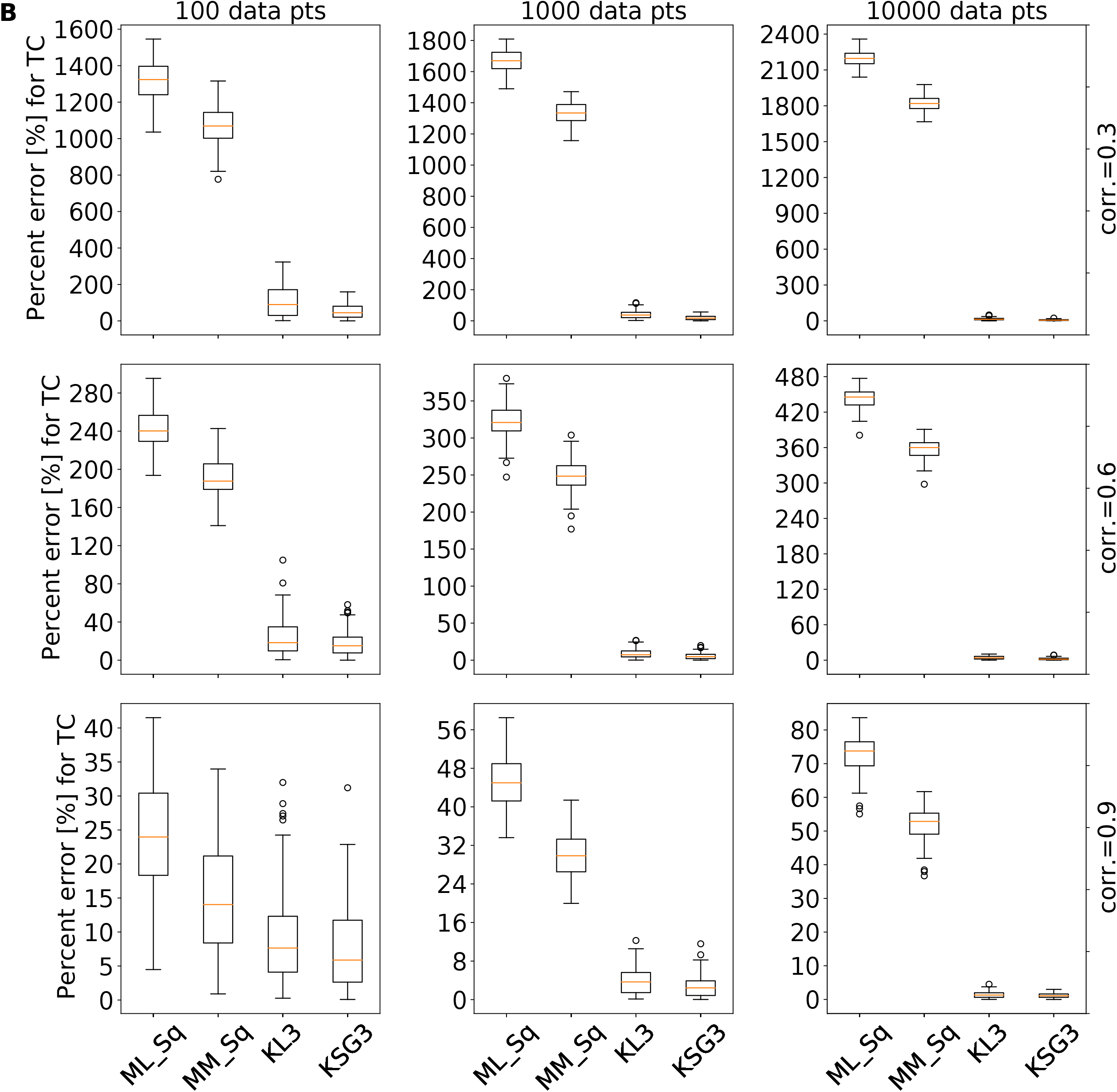
Percent error of different mutual information estimators for multivariate gaussian distribution. Each boxplot represents 100 replicates, with columns representing sample size = {100,1K,10K}, and rows the correlation = {0.3,0.6,0.9}. (A) Percent error (y-axis) for two-way mutual information (MI2) was compared for 3 different methods: ML_Sq=Maximum Likelihood (Shannon’s MI) with fixed width binning (number of bins is determined by square-root), MM_Sq=Miller-Madow formula for MI with square-root for the number of bins, KSG3 =KSG formula for kNN-MI with k=3; (B) same methods compared for total correlation (TC).

**Table 1:**
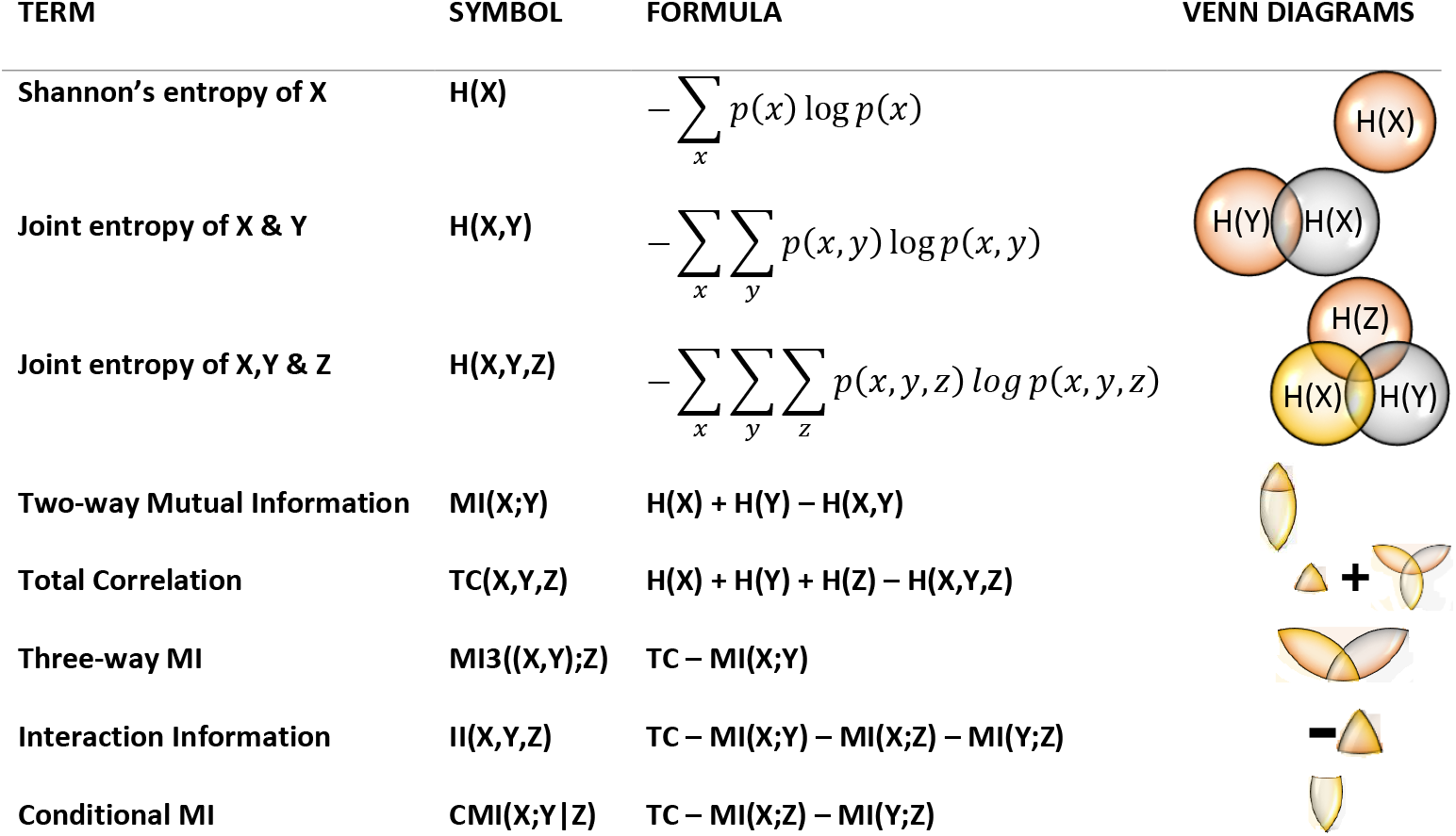
Mutual Information formalism

While two-way MI estimators were studied extensively [22, 26], to our knowledge, no benchmark was done on MI with three or more variables. We repeated the same methodology described above but this time for the 3d Total Correlation (TC) (**Fig. 2B,** Additional file 2: **Fig. S1C-D**). Similar to the 2d case, kNN-based MI estimators KL3 and KSG3 outperformed the other methods. We also examined the other three-way MI quantities, three-way MI (MI3), Interaction Information (II), Conditional Mutual Information (CMI) (see Additional file 2: **Fig. S2-4**) and obtained similar results. We also explored whether a higher kNN value, for example k=10, further improved accuracy. We found that a higher k value (k=10) does not improve the accuracy dramatically compared to that in k=3 (Additional file 2: **Fig. S5,6**), but it did reduce the variance for small correlations (r=0.3).

### *In Silico* GRN Inference performance enhancement

Next, we aim to investigate whether the high precision of MI estimation based on kNN for bi-bin number other thanand tri-variate Gaussian distributions also translates to a high performance in inferring GRN structure compared to other MI estimation methods described above.

To compare the performance of different MI estimators and inference algorithms, we used a total of 15 different synthetic networks: ten synthetic networks from the DREAM3 (Dialogue for Reverse Engineering Assessments and Methods) competition [27] with 50 and 100 genes, respectively, and five networks from DREAM4 with 100 genes. The networks were extracted from documented regulation databases of *E. coli* and *S. cerevisiae* [28]. We used the software GeneNetWeaver 3.1.2b [29] with default settings to generate simulated expression data for each network and performed ten replicates to include the variance in expression data due to stochastic molecular noise. Furthermore, to comply with the majority of available experimental data, we only used the simulated steady state data (Wild type, knockouts, dual-knockouts, knockdowns, multifactorial perturbation) accumulating to 170, 169 and 201 conditions in the 50 gene synthetic networks for *E.coli* 1, *E.coli* 2 and Yeast1/2/3 respectively, 341, 322 and 401 conditions in the DREAM3 100 gene synthetic networks for *E.coli* 1, *E.coli* 2 and Yeast1/2/3 respectively, and 393, 401 conditions in the DREAM4 100 gene networks. We then ran the expression data through our custom Python 3.8 code pipeline to calculate the area under precision-recall curve (AUPR) for each replicate.

In **Fig. 3** we show sorted boxplots of the AUPR values (*y*-axis) comparing six combinations of three inference algorithms (Relevance Networks, RL; Context-Likelihood-Relatedness, CLR; and our Conditional-Mutual-Information-Augmentation, CMIA) and two MI estimators (ML, fixed bin-based; KSG, kNN-based), for five networks with 50 genes (**Fig. 3A**), five networks of 100 genes from DREAM3 (**Fig. 3B**), and five networks of 100 genes from DREAM4 (**Fig. 3C**). In all cases, the kNN-based KSG as the MI estimator improves the performance of the inference algorithms. The improvement is more significant for CMIA, which uses three-way MI calculations, and corroborate the higher percent error we found when estimating TC (**Fig. 2B**).

**Figure 3.**
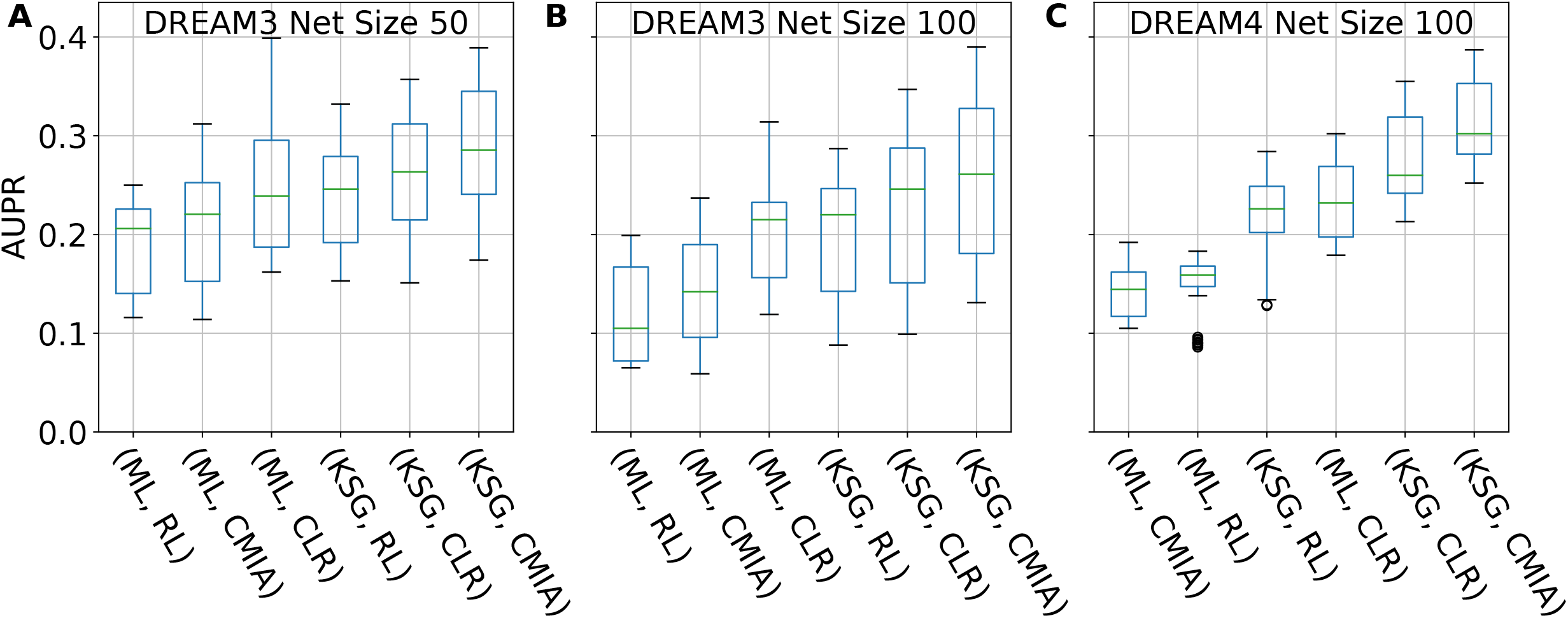
AUPR values for different combinations of MI estimator (ML or KSG) and GRN inference algorithm (RL, CLR or CMIA). (A): Sorted boxplots showing networks of size 50 from DREAM3, (B): Networks of size 100 from DREAM3, (C): Networks of size 100 from DREAM4. For the different network sizes each boxplot represents 50 networks (5 different networks X 10 replicates).

### *In Silico* GRN Inference performance comparison

To verify whether the performance enhancement introduced by kNN-based MI estimators is general for other GRN inference algorithms, we further extended our benchmark to twenty-four different combinations of the four MI estimators (discrete bin-based ML and MM, and kNN-based KL, and KSG) with six inference algorithms described in the Methods section (RL, CLR, ARACNE, SA-CLR, CMIA, CMI2rt) and compared them to the field gold standard combination {(ML, CLR)} (**Fig. 4**). To compare the performance differences quantitatively, we calculated the change in AUPR for each replicate relative to the field’s gold standard combination of CLR inference algorithm with ML for MI calculations. In **Fig. 4** we show the top nine combinations, omitting ARACNE and CMI2rt among the inference algorithms, and KL from the MI estimators because of their poor performance. We also omitted SA-CLR due to its similarity to CLR and CMIA (see full data in Additional file 3: Table S1). The combination of {KSG,CMIA} gave the best median score in the combined networks inspected under each category. It showed a median improvement of 16% and 24% for networks of 50 and 100 genes from DREAM3, respectively (**Fig. 4A, B**), and 34% improvements for networks of 100 genes from DREAM4 (**Fig. 4C**). Furthermore, replacing the MI estimator from ML to KSG in the case of the gold standard ({ML,CLR}} can lead to significant improvement in GRN reconstruction performance, with median increase in AUPR of 8-18%.

**Figure 4.**
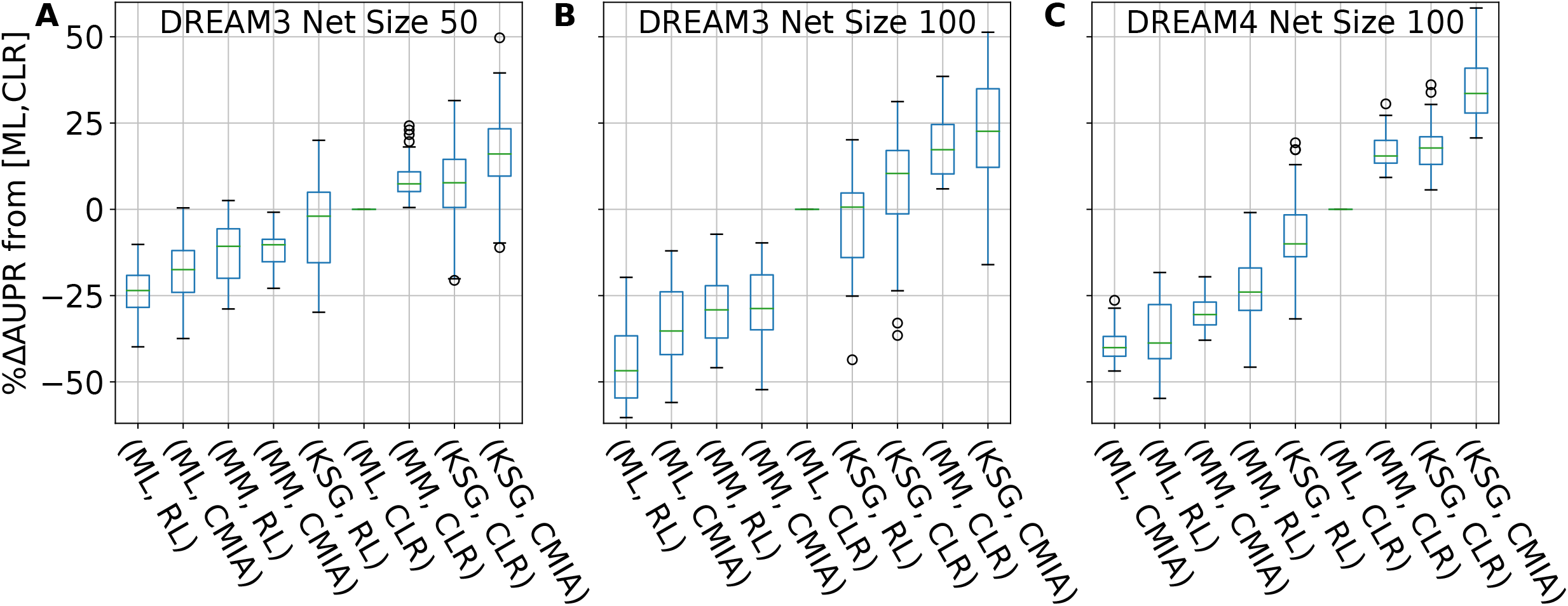
AUPR difference of combinations of MI estimators and inference algorithms relative to the gold standard {ML,CLR]. (A): Sorted boxplots showing comparison for Network size of 50 from DREAM3, (B): and size of 100 from DREAM3, (C): size of 100 from DREAM4. Each boxplot represents 50 networks (5 different networks X 10 replicates). A complete list of tested GRN inference algo & MI estimators can be found in Additional file 2 Table S1

### *In Silico* GRN Inference performance of different organisms

Next, we examined the performance of these different algorithms with regards to different biological organisms, as *E. coli* and *S. cerevisiae* have distinct distributions of different network motifs (Additional file 2: **Fig. S7**), which may lead to different performance in network inference. For example, the fan-out motif, where one gene regulates two (or more) target genes, is more abundant in *E. coli,* while the cascade motif, where a gene regulates a second gene that in turn regulates a third gene, is more abundant in *S. cerevisiae* [7, 30]. In both cases, the three participating genes exhibit some degree of correlation, yet not all are directly connected. The 10 networks from DREAM3 were divided into four *E. coli* networks (**Fig. 5A, C-F**) and six *S. cerevisiae* networks (**Fig. 5B,** Additional file 2: **Fig. S8**). For the combined *E. coli* networks (**Fig. 5A**), KSG greatly improved the performance of both RL and CMIA algorithms but showed only a modest 6% improvement in performance for CLR. For the combined *E. coli* networks, {KSG,CMIA} achieved a median improvement of 20%, but was second best to {MM,CLR}. The performance comparison of the individual *E. coli* networks (**Fig. 5C-F**) showed that {KSG,CMIA} was the best performer on three out of four networks. Furthermore, replacing ML with KSG when combined with CLR improved the performance by 10-15% except in the case of DREAM3 Ecoli2-Size100 (**Fig. 5F**). In the *S. cerevisiae* networks, again KSG improved all algorithms, and most significantly CMIA, and showed a median improvement of 18%. Several replicates did not show any performance improvement, indicating the significance of stochasticity even though all kinetic parameters for each network were identical.

**Figure 5.**
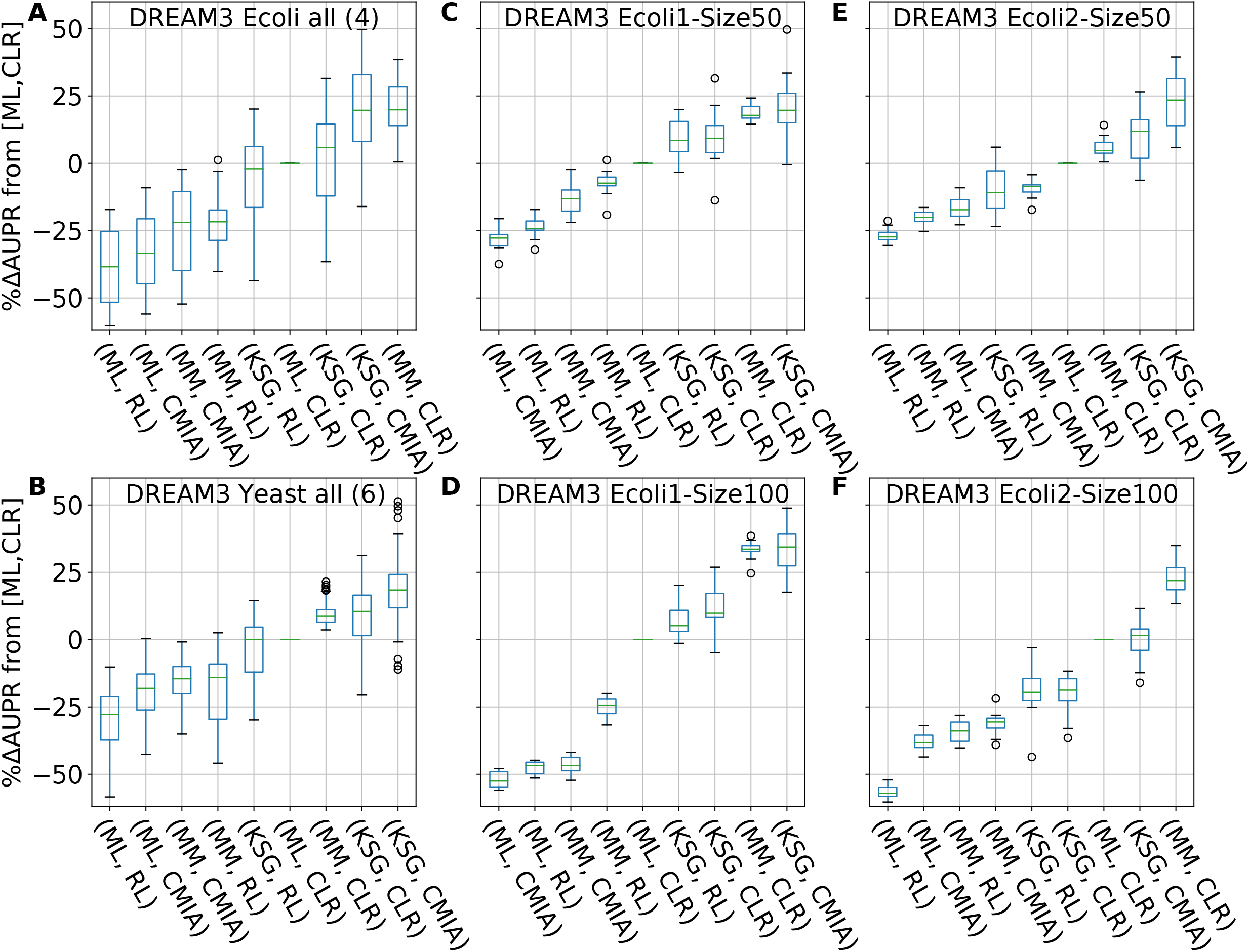
Performance comparison of GRN reconstruction for different in silico networks modeled from E. coli & Yeast. x-axis shows different combinations of [MI estimator, inference algo], y-axis shows percentage AUPR difference (increase or decrease) relative to the gold standard combination [ML,CLR]. (A): Sorted boxplots of the combined four E.coli networks from DREAM3. Each boxplot represents 40 networks (4 different networks X 10 replicates). (B) same as (A) but for the six Yeast networks. (C)-(F): Sorted boxplots of the 4 different E.coli networks from DREAM3. Each boxplot represents 10 replicates. A complete list of tested MI estimators & GRN inference algo can be found in Additional file 2 Table S2

In summary, out of 24 combinations of MI estimators and inference algorithms, the combination {KSG,CMIA} yielded the best median score in 13 out of the 15 networks inspected (except networks DREAM3 Yeast1-Size50 & Ecoli2-Size100, **Fig. 5C-F,** Additional file 2: **Fig. S8 and S9**). Therefore, we conclude that using kNN-based KSG to calculate MI improved the performances of the inference algorithms evaluated in most cases.

### Computational cost

Computational cost is a major concern when applying kNN-based methods. We measured the time required to calculate all the two- and three-way interactions in a 50 gene network (1125 pairs and 19600 triplets, respectively, after taking symmetry into account) with different data size [100, 250, 500, 1000] for three MI estimation methods: FB-ML, kNN-KL and kNN-KSG. The code for the three estimators was written in Python 3.8, used built-in functions from Numpy v1.19 and Scipy v1.5, and was run on a single core of a desktop [Intel Xeon E5-1620 @ 3.6 GHz]. As seen in **Fig. 6** FB-ML was the fastest, as histogram-type calculations have been optimized in Python over the years. FB-ML was also insensitive to data size (in the tested range). While the python-based KSG implementation was most computationally heavy, the time was tractable (under 400 s even for the largest data size (1000) and 3d calculation). The speed could be further boosted by rewriting the code in C/C++, similar to what was done by Meyer et al. [31] and Sales et al. [24]. Furthermore, the KD-Tree class of algorithms [32], which was in the main core of this work’s implementation, could greatly benefit from multiple cores or parallel processing. After building the initial tree, distance calculation between neighbors can proceed in parallel, offering 4-to-16 fold improvement in speed on a current personal computer, depending on the number of available cores.

**Figure 6.**
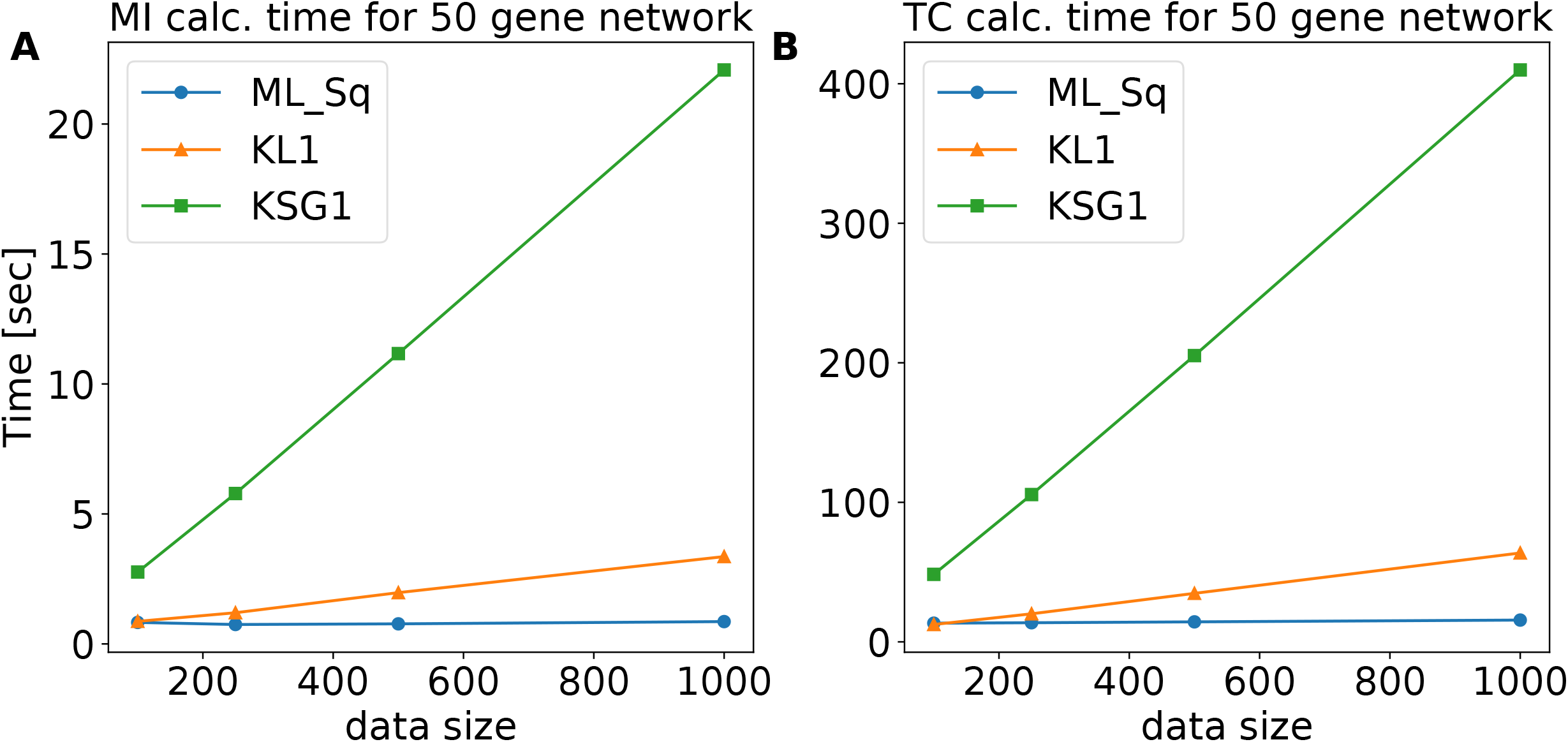
Computation time vs. different data sizes for a network of 50 genes. (A) The calculation is performed over 1125 pairs for data sizes of [100, 250, 500, 1000]. (B) The calculation is performed over 19600 triplets for data sizes as in the left panel

## Discussion

To date, a plethora of discretization methods, MI estimators, and inference algorithms exist in the literature to reconstruct GRNs. Some common methods are available in the R/Bioconductor package *minet* [31] and in Julia language [33]. In fact, as different methods have certain advantages depending on the investigated scenario and constraints, it is advantageous to consider and compare the performance of different combinations of multiple methods [34].

### kNN-based MI estimator for date discretization/density estimation outperforms fixed-bin-based estimations

Here, we demonstrate that the MI estimator KSG based on kNN yields smaller errors compared to other MI estimation methods using discretized fixed bins in the case of a bi- and tri-variate Gaussian distribution. KSG proves to be robust against different data sizes and correlations as well as the *k* parameter used, unlike FB methods where the parameter used (number of bins) has a large effect on accuracy of the MI estimator. In principle, one can achieve smaller errors using MI based on discretized bins by choosing a different bin number other than the rule of thumb 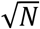, for correlations smaller than 0.9. However, *a priori* one does not know the correlation strength. In fact, estimating the correlation strength is what one tries to achieve when using MI. We also note that the gene expression profiles of different synthetic networks and real experimental systems could be better described by distributions other than Gaussian. Fortunately, the analytical solution to the mutual information of a few of these distributions can be calculated [35] and will be explored in future work.

Note that in this work we did not compare the performance of another frequently used binning method, adaptive partitioning, which is computationally faster than kNN for large data sets. In brief, adaptive partitioning is a general term referring to three methods that divide the data uniformly between the bins. The first method is equal frequency in which the bin size varies to allow for equal number of data points in each bin. The second method is equiprobable partitioning [21, in which data is ranked and partitioned in the middle, and Pearson chi-square test is used to determine the number of sub-partitions, where the significance level of the chi-square test can be tuned (1%-5%) according to the size of the data. This method works well for 1d data, but it has some ambiguity when implemented in higher dimensions in that data points must be ranked according to one of the axes (or more in >2d), and there are no appropriate rules to rank multidimensional data points. The third method is Bayesian blocks [36], which uses a Bayesian statistics approach to attempt to find the optimal number of bins and their sizes by maximizing a fitness function that depends on those two parameters. While this is a seemingly promising approach, it is unclear how to implement such a method beyond 1D. Because of these reasons, we did not include this binning method in the comparison.

Another previously used method in the literature is KDE [14], but it is the most computationally costly and requires large data sets. It approximates the data distribution using a predefined known distribution (i.e., a Gaussian) with user-defined smoothing parameters. This practice can be problematic because in most cases the underlying data distribution is unknown, and experimental data is much sparser than required to achieve results similar to other, simpler methods, such as FB.

### kNN-based MI estimator KSG in combination with CMIA achieves the highest accuracy but may subject to data stochasticity

It is clear from **Fig. 3** and **4** that the combination of kSG-based MI estimation and inference algorithm CMIA achieved the highest precision and recall when reconstructing an unknown network. Yet, this combination also showed a large variation in the performance enhancement. As shown in **Fig. 4, 5A-B**, we observed that when KSG was combined with CLR or CMIA, a few replicates did not show any performance improvement, or even had a decreased performance indicated by the negative %ΔAUPR value, as indicated by the bottom whisker of the boxplot.

To investigate the source of this variation in the ensemble network plots we inspected different combinations of MI estimators, inference algorithms, data size used, and individual networks (**Fig. 5C-F,** Additional file 2: **Fig. S8**). We found that higher k values (up to k – 15) did not affect the variability in the AUPR results (Additional file 2: **Fig. S10**). However, MI calculation done by KSG exhibited large variations in performance when smaller data size was used as that in the case of 50 gene networks. For example, in **Fig. 5C,E**, KSG showed a performance enhancement in the range of ~ 25-35% for the three different inference algorithms, but the variability was reduced by half when ML instead of KSG was used. This was also shown in the large variance calculated for KSG for a Gaussian distribution (Additional file 2: **Fig. S1D, left column**). This observation indicates that KSG is more sensitive to stochasticity (intrinsic noise) when data size is smaller than a few hundred points. Our choice of algorithm KSG-1 over KSG-2 (see methods) was intended to keep a low statistical error and thus, low variability. However, using total correlation and two-way mutual information to calculate other measures, such as interaction information (**Table 1**), can lead to higher errors as the systematic errors might not cancel out as we have demonstrated in this work. Additionally, when using KSG, we set negative values of total correlation and two-way mutual information to zero (due to statistical fluctuations at low correlation values) prior to calculating the other 3d MI quantities. This practice does not change the results for pairs or triplets with highly positive MI values, but in some cases could lead to increased errors as gene pairs with low MI would be ranked differently.

We note that two networks (DREAM3 Yeast1-Size50 & *E.*coli2-Size100) out of the 15 networks investigated showed no performance enhancement when using {KSG, CMIA} compared to the Gold Standard {ML, CLR} (**Fig. 5F,** Additional file 2: **Fig. S8)**. It is unclear why the performance did not improve in these two cases based on the largely similar statistics of different motifs of the ten networks from DREAM3 (Additional file 3: **Table S3**). It could be due to a specific sub-structure of this network, but further analysis is needed.

Another important result we observed (**Fig. 4, 5**) is that the combination {MM,CLR} achieved higher AUPR for all replicates over {ML,CLR}. This is probably due to the size of the data used, as MM was developed to correct the bias in MI estimation for small data sets. We thus suggest using this combination as the new gold standard of the field when working with similar data sizes and when fixed-binning for data discretization is preferred.

## Conclusions

In summary, we have shown that the kNN-based KSG MI estimator improves the performance of inference algorithms, especially ones that use three-way MI calculations. This result corroborates our observations in comparing MI calculations against the analytical solution of two-way MI of a bi-variate Gaussian distribution and the total correlation of a tri-variate Gaussian distribution. Furthermore, the combination of CMIA and KSG give the overall best performance, and hence should be preferred when precision and recall are more important than speed when reconstructing a GRN. Looking forward, the goal of complete reconstruction of GRNs may require new inference algorithms and probably MI in more than three dimensions.

## Methods

### Calculate mutual information of multiple variables

In **Table 1**, we summarize the formalism for calculating MI. Shannon’s entropy is the basic building block of MI and represents the randomness of a variable: the more random it is, the more uniformly it is distributed, which gives a higher entropy. For our purposes, X, Y, or Z is a vector (*x_1_, x_2_, …, x_n_*), (*y_1_, y_2_, …, y_n_*) or (*z_1_, z_2_, …, z_n_*) representing a specific gene’s expression profile (data *x, y or z*) under different conditions/perturbations (*n* steady-states) or as a function of time (*n* time points). Two-way MI is defined as the shared (or redundant) information between the two variables X and Y {Table 1} and can be visualized by a Venn diagram {**Table 1 right column**}.

While MI for two variables (genes or dimensions) is readily understood, for three variables or more, new measures arise including Total correlation (TC), Three-way MI (MI3), Interaction Information (II) and Conditional MI (CMI) (**Table 1**). Unfortunately, the term ‘three-way MI’ has been used loosely in the literature to refer to all four of these measures, and because they represent distinct aspects of statistical dependence, in the context of GRN reconstruction, this can lead to different realizations. Unlike other MI quantities, Interaction-Information is hard to visualize using a Venn diagram, as it can have both positive and negative values. It is common to regard negative II as “Redundancy”, the shared information between all variables, and positive II as “Synergy”. Synergy can be interpreted as new information gained on the dependence between two variables {X,Y} when considering the contribution of a third variable {Z} on either {X} or {Y} *v.s.* without considering it, or mathematically: II=CMI(X;Y|Z)-MI(X;Y).

To calculate the marginal and joint entropies of two variables (X and Y), we first need to know the probability of each data point. For discrete data, we can approach the underlying probability *p(x)* by calculating the frequency 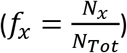 where N_x_ is the number of data points with value *x*, and N_Tot_ is the total sample size. For the continuous data case, the calculation is more complex. Although Shannon extended his theory for continuous data by replacing the summation with integrals, it is common practice in the field to discretize the data first so one can work with the discrete formalism (**Table 1**). The simplest discretization method is to use **fixed (width) binning** (**FB**) (**Fig. 1A**), but the optimal binning choice depends on the shape of the distribution and data size. For normally distributed data, the rule of thumb is to use the square-root of the data size as the number of bins.

### k-nearest-neighbor (kNN)

Other than evaluating the probability densities to calculate mutual information, Kozachenko and Leonenko (KL) calculated the marginal and joint entropies (and the MI by summation) from the mean distance to the kth-nearest neighbor [25]. To minimize errors when combining entropies of different dimensions, Kraskov et al. calculate the MI directly [22]. KSG developed two algorithms, I^(1)^ and I^(2)^ (hereafter, KSG-1 and KSG-2), to minimize errors when estimating MI compared to previous methods. We chose KSG-1 (defined below as MI_KSG) as it gives slightly smaller statistical error (dispersion). Note that although KSG-1 gives relatively larger systematic errors than KSG-2, these systematic errors do not change the ranking of the output values (from high to low), which is what we use in downstream analysis. An additional note is that using kNN can lead to negative values for mutual information, which contradicts Shannon’s theorem. Negative values are caused by statistic fluctuations when there is no correlation between variables. Therefore, in such a situation, we set negative values to zero (except for Interaction Information, where it is meaningful). To calculate MI using the KSG method, we use the following formulas:

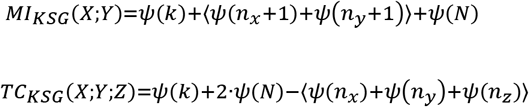

Where ψ(x) is the digamma function, N is the number of data points, ni is the number of points xj whose distance from xi is less than ε(i)/2, and ε(i)/2 is the distance from u_i_=(xi,yi,zi) to its kth neighbor, as illustrated in Fig. 1(a) of [22]

### *In Silico* GRN Inference comparison

MI calculations are used to infer interactions between genes to reconstruct the underlying GRN structure. To test the performance of different methods, we followed the methodology of the *in silico* network inference challenges of the **Dialogue for Reverse Engineering Assessments and Methods** (DREAM) competitions DREAM3/4 [27] as depicted in **Fig. S11**.

#### 1. Simulating gene expression data

we used GeneNetWeaver [29] to generate steady-state and time-series gene expression datasets for realistic *in silico* networks of sizes of 50, and 100 genes containing various experimental conditions (knockouts, knockdowns, multifactorial perturbation, etc.). GeneNetWeaver uses a thermodynamic model to quantify mRNA transcription and regulation with added molecular and experimental noise.

#### 2. Discretizing/density estimation

To handle the continuous expression data, we chose either:

a. Density estimation by fixed bin. We used the common practice sqrt(n), where n = number of data points (in our case, different experimental conditions), as the number of bins.
b. Density estimation by k-Nearest Neighbor (kNN). We chose k=3 as a good compromise between precision and computation cost as discussed in the previous section.

#### 3. Mutual Information estimation

Depending on our previous selection, we chose between several MI estimators:

a. For the fixed-bin discretizing method, we used either Shannon’s formula (also referred to as *Maximum Likelihood,* ML) or *Miller-Madow* (MM) estimator.
b. For kNN we used either KL or KSG formulas for MI.

#### 4. GRN inference algorithms

We used popular algorithms in the field that use either only two-way MI or both two- and three-way MI to infer undirected network structure by sorting predicted interacting gene pairs from most probable to least probable. Each algorithm starts with a MI matrix containing calculation for all possible pairs (some use all possible triplets) and applies different rules to filter results and sort the gene pairs (see summary below). We used the same MI matrices for a fair comparison between the inference algorithms. The following algorithms were used in our comparison:

a. Relevance Network (RL) – Gene pairs are sorted according to their MI(X;Y) value from highest to lowest, and a threshold applied to truncate non-significant results [10]. We didn’t set a threshold to maximize AUPR (see below).
b. Algorithm for the Reconstruction of Accurate Cellular Networks (ARACNE) – Same as RL with the addition of <u>Data Processing Inequality</u> (DPI), which means for every three genes MI is calculated for each pair and the pair with the lowest MI is removed if the difference is larger than some threshold [11]. In our implementation, we set the threshold to zero, so we always removed the lowest interacting pair. On the other extreme, where we kept all the pairs, ARACNE is the same as RL.
c. Context Likelihood of Relatedness (CLR) – Background correction is performed by calculating Z-score for the MI of each gene interacting with all other genes, and then gene pairs are sorted by their mutual Z-score [12]. We didn’t use B-spline smoothing in the density estimation step in accordance with the implementation in the R-package *Minet* [31].
d. Synergy Augmented CLR (SA-CLR) – Same as CLR, with the difference that now the highest Interaction-Information term is added to MI prior to performing the background correction [17]
e. Conditional Mutual Information Augmentation (CMIA) – Similar to SA-CLR but we used conditional mutual information instead of interaction-information.
f. Luo et al. MI3 (hereafter CMI2rt) – We assumed two regulators for each target gene, and for each target gene we searched for the best {R1,R2} pair that maximizes: CMI(T;R1|R2)+CMI(T;R2|R1) [14]

#### 5. GRN performance evaluation

To evaluate the performance of common algorithms in the field, we used known (true) synthetic networks and counted the number of true and false positives (TP and FP respectively) predictions as well as true and false negative (TN and FN respectively) **(Fig. S12).** This allowed us to plot precision (Precision = TP/(TP+FP)) *v. s.* recall (Recall = TP/(TP+FN)) and calculate the area under precision-recall curve (AUPR). As biological networks are sparse on edges, AUPR is considered a better metric than AUROC (area under the receiver operating characteristic curve, which is the false positive rate FPR = FP/(FP+TN v.s. recall) as mentioned elsewhere [37].

## List of abbreviations

GRN: Gene regulatory network
ODE: Ordinary differential equations
MI: Mutual information
PDF: Probability density functions
FB: Fixed (width) binning
AP: Adaptive partitioning
kNN: k-nearest neighbor
KDE: Kernel density estimator
CLR: Context likelihood of relatedness
CMIA: Conditional mutual information augmentation
KSG: Kraskov-Stoögbauer-Grassberger
RL: Relevance networks
ARACNE: Algorithm for the Reconstruction of Accurate Cellular Networks
SA-CLR: Synergy-Augmented CLR
ML: Maximum likelihood
MM: Miller-Madow
KL: Kozachenko-Leonenko
TC: Total correlation
MI3: Three-way MI
II: Interaction information
CMI: Conditional mutual information
DREAM: Dialogue for reverse engineering assessments and methods
AUPR: Area under precision-recall curve
CMI2rt: Luo et al. inference algorithm named MI3
DPI: Data Processing Inequality

## Declarations

### Availability of data and materials

The software GeneNetWeaver used to generate the datasets in the current study is available in the GitHub repository, https://github.com/tschaffter/genenetweaver

The code and scripts used for analysis and to generate the plots in the current study are available in the GitHub repository, https://github.com/XiaoLabJHU/GRN_Inference

The GRN inference pipeline implemented here is modular, one can use specific functions to calculate MI quantities based on kNN and integrate the output matrix into a different inference algorithm than the ones implemented in this work.

### Authors’ contributions

LS, ER and JX have conceived the study. LS implemented the code to analyze and interpret the data. JX and PC contributed to the interpretation of data. LS have drafted the work. JX and PC have substantially revised the manuscript. All authors read and approved the final manuscript.

## Acknowledgements

Basilio Cieza Huaman for discussion and comments and Miriam Asnes for providing technical writing review.

## Supplementary information

**Additional file 1: Supplementary information Appendix 1-2**

**Appendix S1**: Analytical solution for a multivariate Gaussian distribution

**Appendix S2**: Miller-Madow correction to Shannon’s entropy

**Additional file 2: Supplementary figures S1-10**

**Figure S1**: 100 replicates of two-way mutual information (MI2) & total correlation (TC) for multivariate gaussian dist. With sample size = {100,1K,10K}, correlation = {0.3,0.6,0.9}.

**Figure S2-S4**: boxplots of percent error of three different mutual information estimators for 100 replicates of tri-variate gaussian dist.

**Figure S5**: boxplots of percent error of two-way mutual information calculated based on kNN methods for 100 replicates of bi-variate gaussian dist. With sample size = {100,1K,10K}, correlation = {0.3,0.6,0.9}.

**Figure S6**: boxplots of percent error of Total Correlation calculated based on kNN methods for 100 replicates of tri-variate gaussian dist.

**Figure S7**: Common 3-node network motifs

**Figure S8:** *Sorted boxplots of* percentage AUPR difference (increase or decrease) relative to the gold standard combination [ML,CLR] *for different combinations of MI estimator and GRN inference algorithm for the 6 different Yeast networks from DREAM3.*

**Figure S9:** *Sorted boxplots of* percentage AUPR difference (increase or decrease) relative to the gold standard combination [ML,CLR] *for different combinations of MI estimator and GRN inference algorithm for the 5 different networks of 100 genes from DREAM4.*

**Figure S10:** Area Under Precision-Recall curve (AUPR) vs. different number of bins or k-neighbors.

**Figure S11**: The different steps for evaluating GRN inference performance.

**Figure S12:** A schematic GRN inference example. The true network contains 10 genes (a.k.a. nodes), and 11 interactions (or edges). The prediction algorithm correctly predicted 6 times (True positive), missed 5 interactions (False negative), and predicted 2 interactions that did not exist (False positive).

**Additional file 3: Supplementary information table S1-3**

**Table S1**: Median AUPR values for different combinations of MI estimator and GRN inference algorithm for different network sizes

**Table S2**: Median AUPR values for different combinations of MI estimator and GRN inference algorithm for different organisms

**Table S3**: Characteristics of the 10 synthetic networks from DREAM3 and statistics of the different 3-node network motifs extracted.

