## Additional file 1 for "Gene regulation network inference using k-nearest neighbor-based mutual information estimation-Revisiting an old DREAM"

**Additional file 1: Supplementary information Appendix 1-2**

**Appendix S1**: Analytical solution for a multivariate Gaussian distribution

Shannon [1] showed that the entropy term of a multivariate Gaussian distribution is given by:

$$H\left( X \right)=\frac{1}{2}\log\left[ \left( 2\pi e \right)^{n}\left| COV \right| \right]$$

Where |COV| represents the covariance matrix. For simplicity we set all the correlations between variables to be equal to ρ.

As all MI quantities can be calculated by their entropy components, we have:

$$MI\left( X;Y \right)=H\left( X \right)+H\left( Y \right)-H\left( X,Y \right)=-\frac{1}{2}\log(1-\rho^{2})$$

$$TC\left( X;Y;Z \right)=H\left( X \right)+H\left( Y \right)+H(Z)-H\left( X,Y,Z \right)=-\frac{1}{2}\log(1-3\cdot\rho^{2}+2\cdot\rho^{3})$$

**Appendix S2**: Miller-Madow correction to Shannon’s entropy

Due to the logarithmic nature of Shannon’s entropy:

$$H^{Shan}\left( X \right)=-\sum_{x} p\left( x \right)\log\left( p\left( x \right) \right)$$

Under or overestimating p(x) by the same value gives different errors on the entropy calculation, leading to bias (downwards). Miller and Madow proposed to correct the bias in Shannon’s entropy by adding the asymptotic bias term [2]:

$$H^{MM}=H^{Shan}+\frac{\left\{ non\_empty\_bins \right\}-1}{2N}$$

Where N is equal to the data size.

Two-way mutual information and higher dimension measures can be calculated by summation of entropies, for example in the case of two-way MI:

$${MI}^{MM}\left( X;Y \right)=H^{MM}\left( X \right)+H^{MM}\left( Y \right)+H^{MM}\left( X,Y \right)$$
