## Additional file 2 for "Gene regulation network inference using k-nearest neighbor-based mutual information estimation-Revisiting an old DREAM"

**Additional file 2: Supplementary figures S1-10**

**Figure S1**: 100 replicates of two-way mutual information (MI2) & total correlation (TC) for multivariate gaussian dist. With sample size = {100,1K,10K}, correlation = {0.3,0.6,0.9}. (A) MI2 with natural log base calculated using Maximum Likelihood with fixed width binning (FB), where the shaded area represents mean +/- 2std. (B) MI2 based on KSG k-nearest-neighbor (KNN). (C) TC based on FB. (D) TC based on kNN

C

A

| 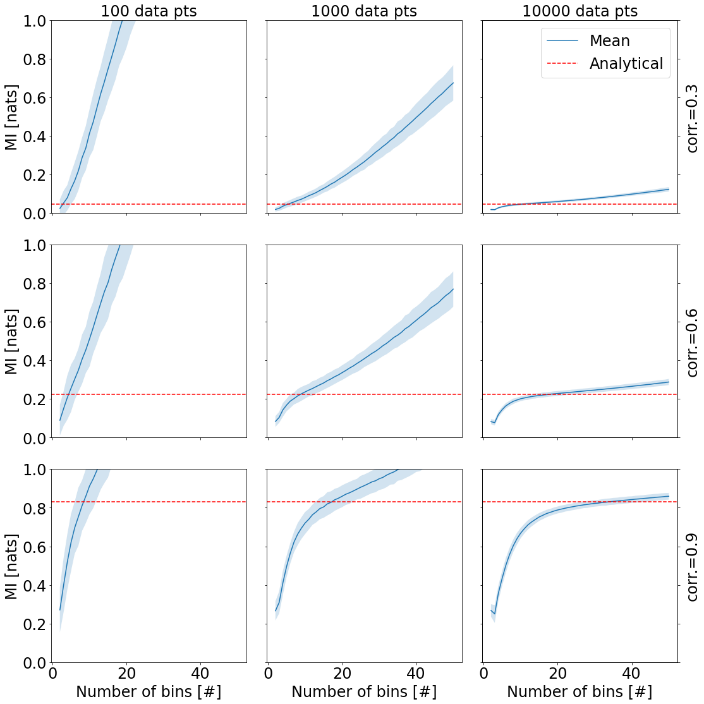  B  D | 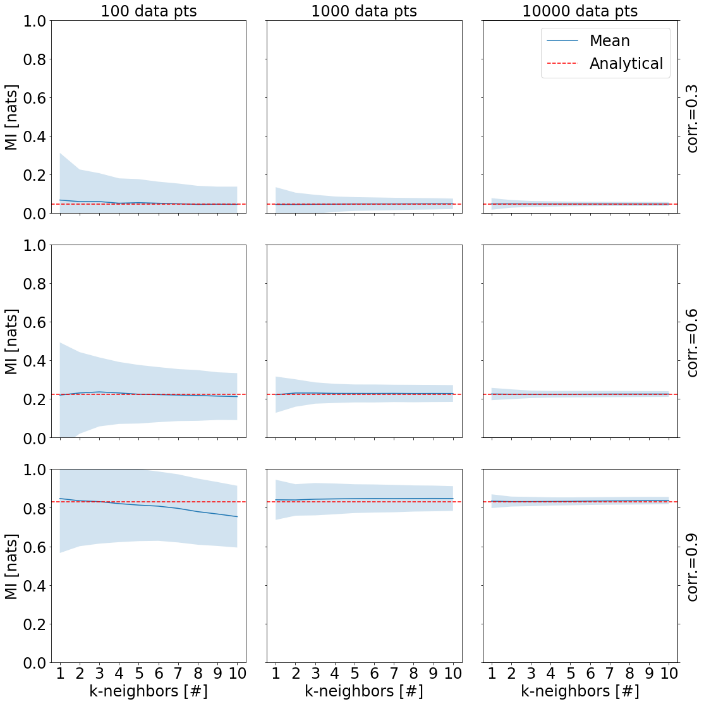 |
| --- | --- |
| 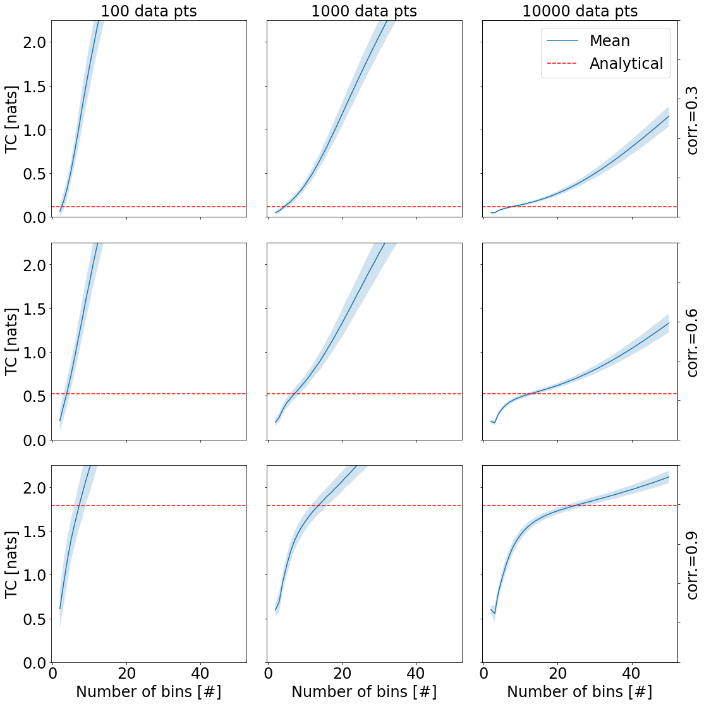 | 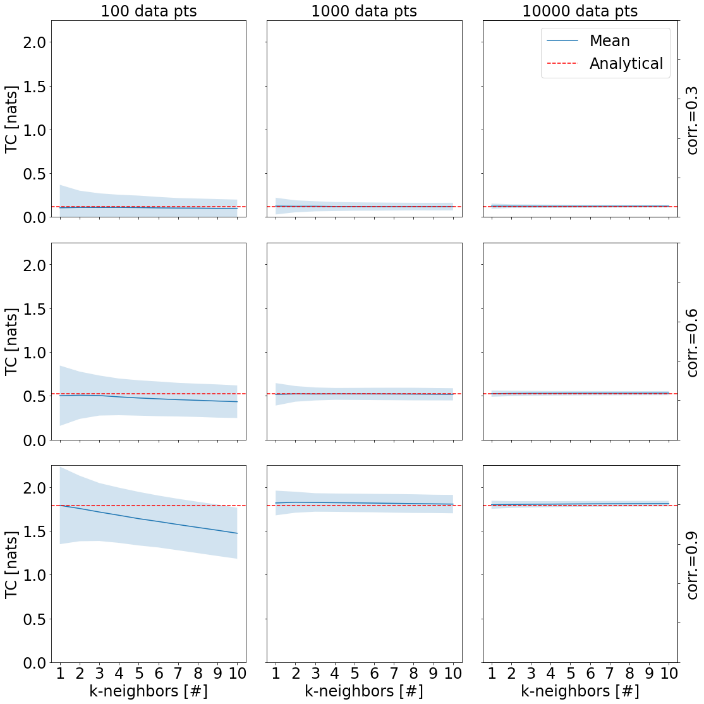 |

**Figure S2**: boxplots of percent error of three different mutual information estimators for 100 replicates of tri-variate gaussian dist. With columns representing sample size = {100,1K,10K}, and rows the correlation = {0.3,0.6,0.9}. 9 subplots showing percent error for Interaction Information (II) for 3 different methods: Sqrt(N)=Shannon’s MI with fixed width binning (number of bins is determined by square-root), MM_Sq=Miller-Madow formula for MI with square-root for the number of bins, kNN3=KSG formula for MI with k=3.
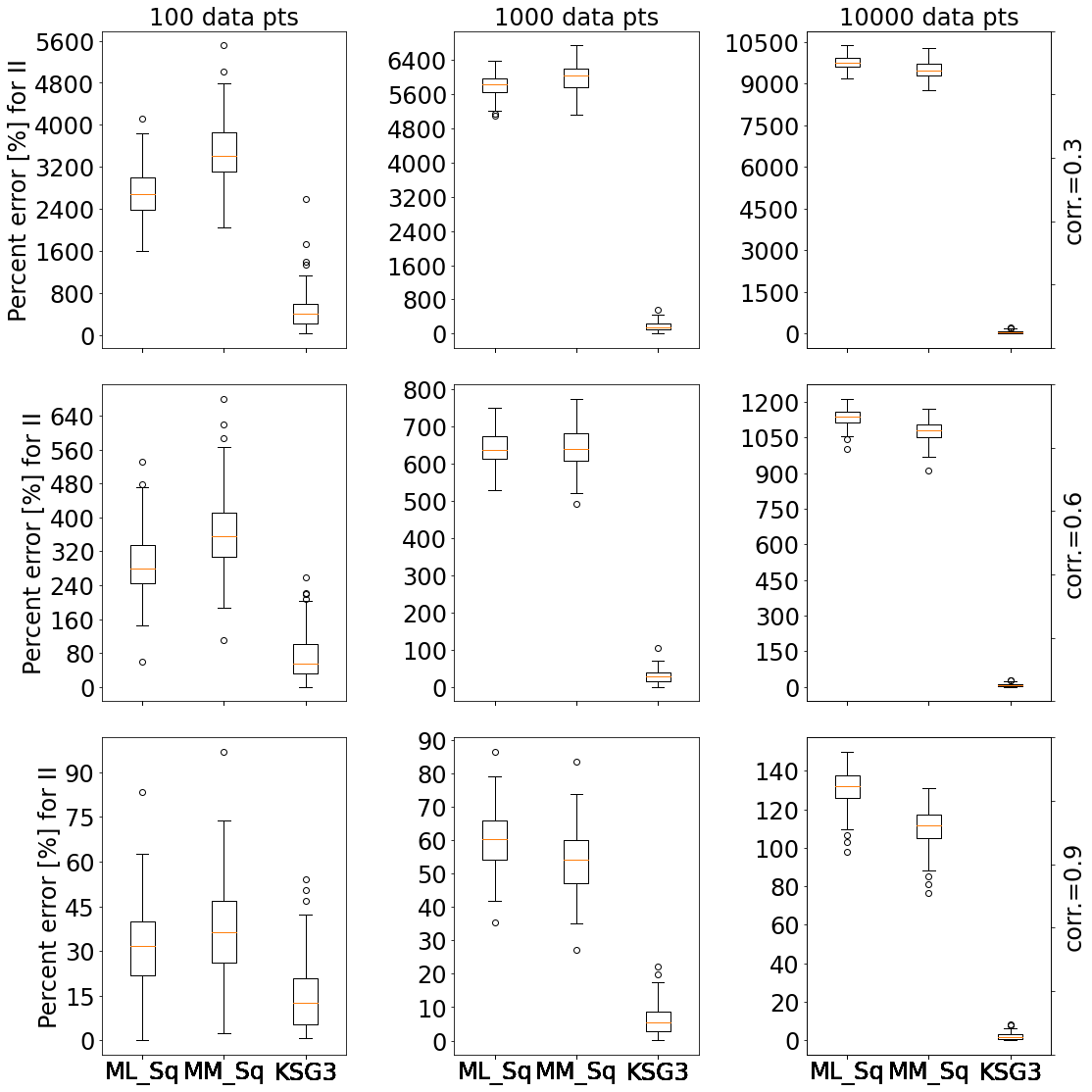

**Figure S3**: boxplots of percent error of three different mutual information estimators for 100 replicates of tri-variate gaussian dist. With columns representing sample size = {100,1K,10K}, and rows the correlation = {0.3,0.6,0.9}. 9 subplots showing percent error for Conditional Mutual Information (CMI) for 3 different methods: Sqrt(N)=Shannon’s MI with fixed width binning (number of bins is determined by square-root), MM_Sq=Miller-Madow formula for MI with square-root for the number of bins, kNN3=KSG formula for MI with k=3.

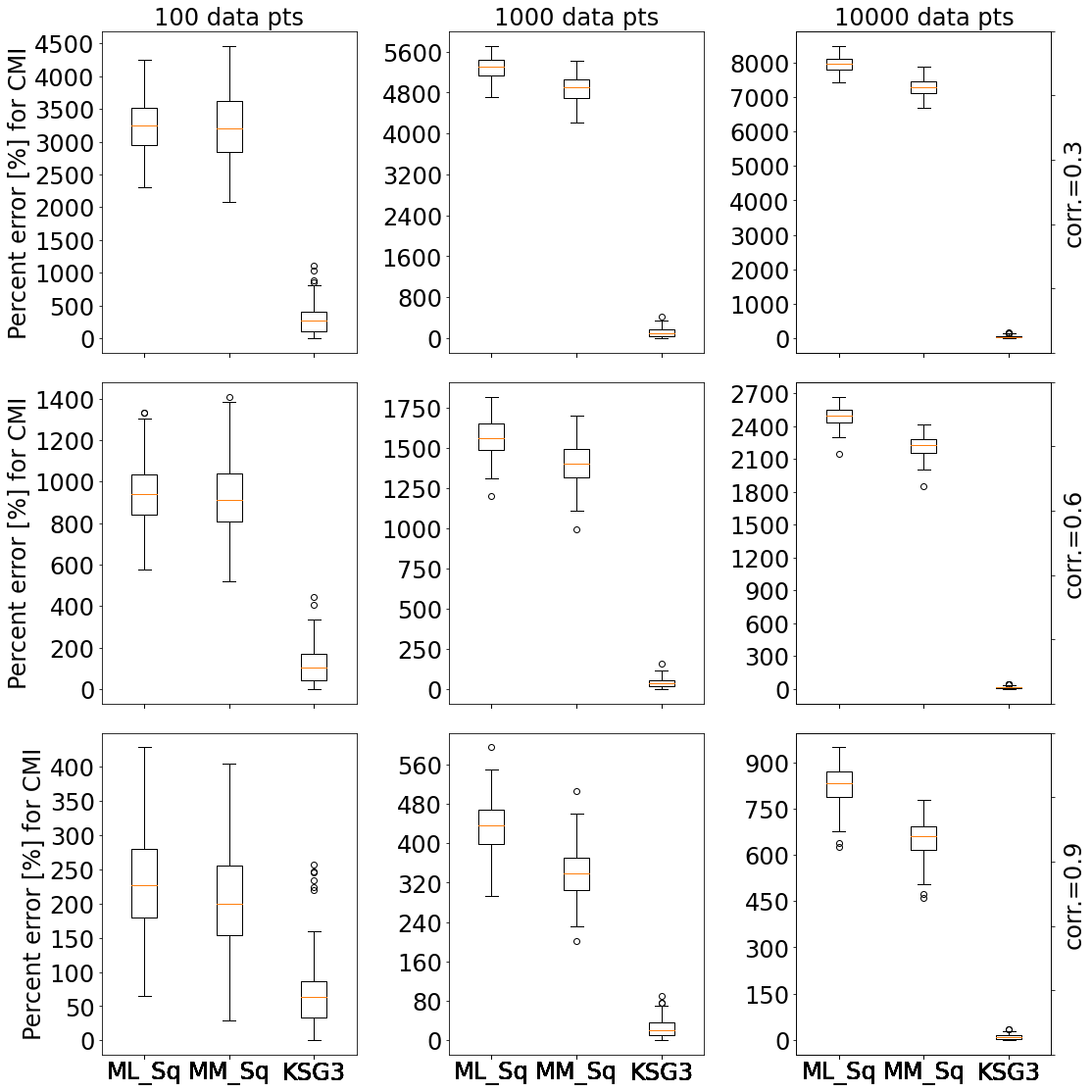

**Figure S4**: boxplots of percent error of three different mutual information estimators for 100 replicates of tri-variate gaussian dist. With columns representing sample size = {100,1K,10K}, and rows the correlation = {0.3,0.6,0.9}. 9 subplots showing percent error for Three-way Mutual Information (MI3) for 3 different methods: Sqrt(N)=Shannon’s MI with fixed width binning (number of bins is determined by square-root), MM_Sq=Miller-Madow formula for MI with square-root for the number of bins, kNN3=KSG formula for MI with k=3.

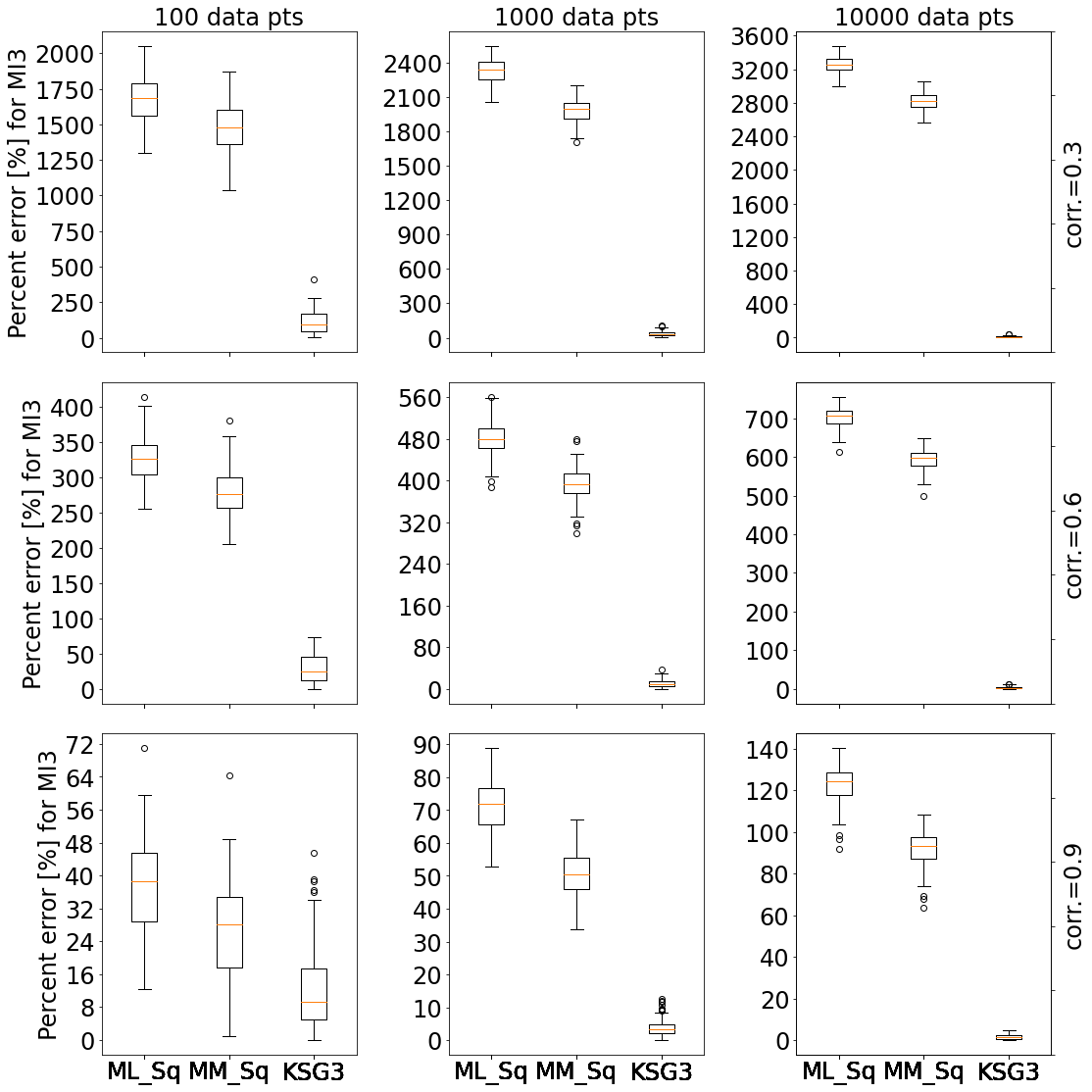

**Figure S5**: boxplots of percent error of two-way mutual information calculated based on kNN methods for 100 replicates of bi-variate gaussian dist. With sample size = {100,1K,10K}, correlation = {0.3,0.6,0.9}. We compare KL and KSG methods for k=1,3,10.

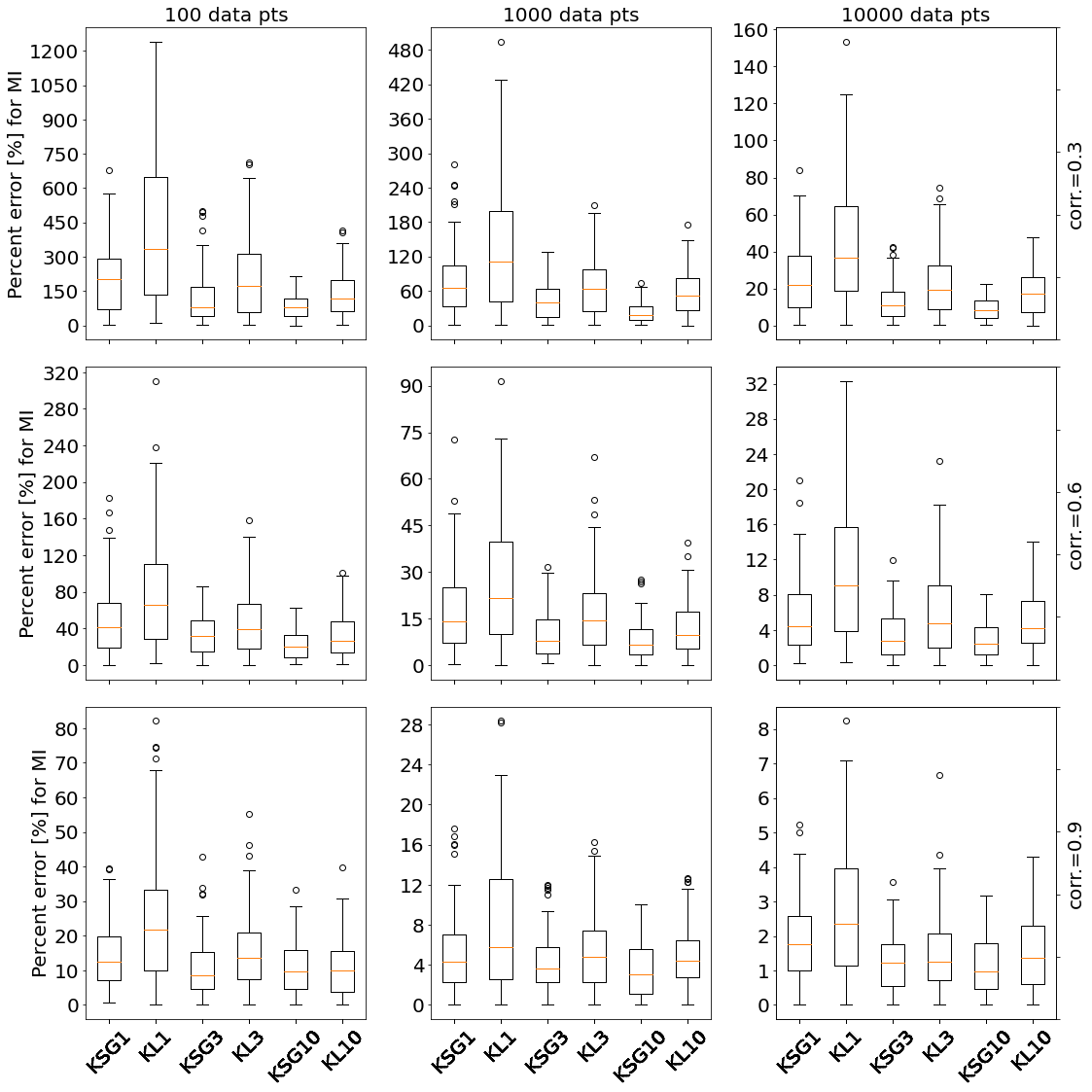

**Figure S6**: boxplots of percent error of Total Correlation calculated based on kNN methods for 100 replicates of tri-variate gaussian dist. With sample size = {100,1K,10K}, correlation = {0.3,0.6,0.9}. We compare KL and KSG methods for k=1,3,10.
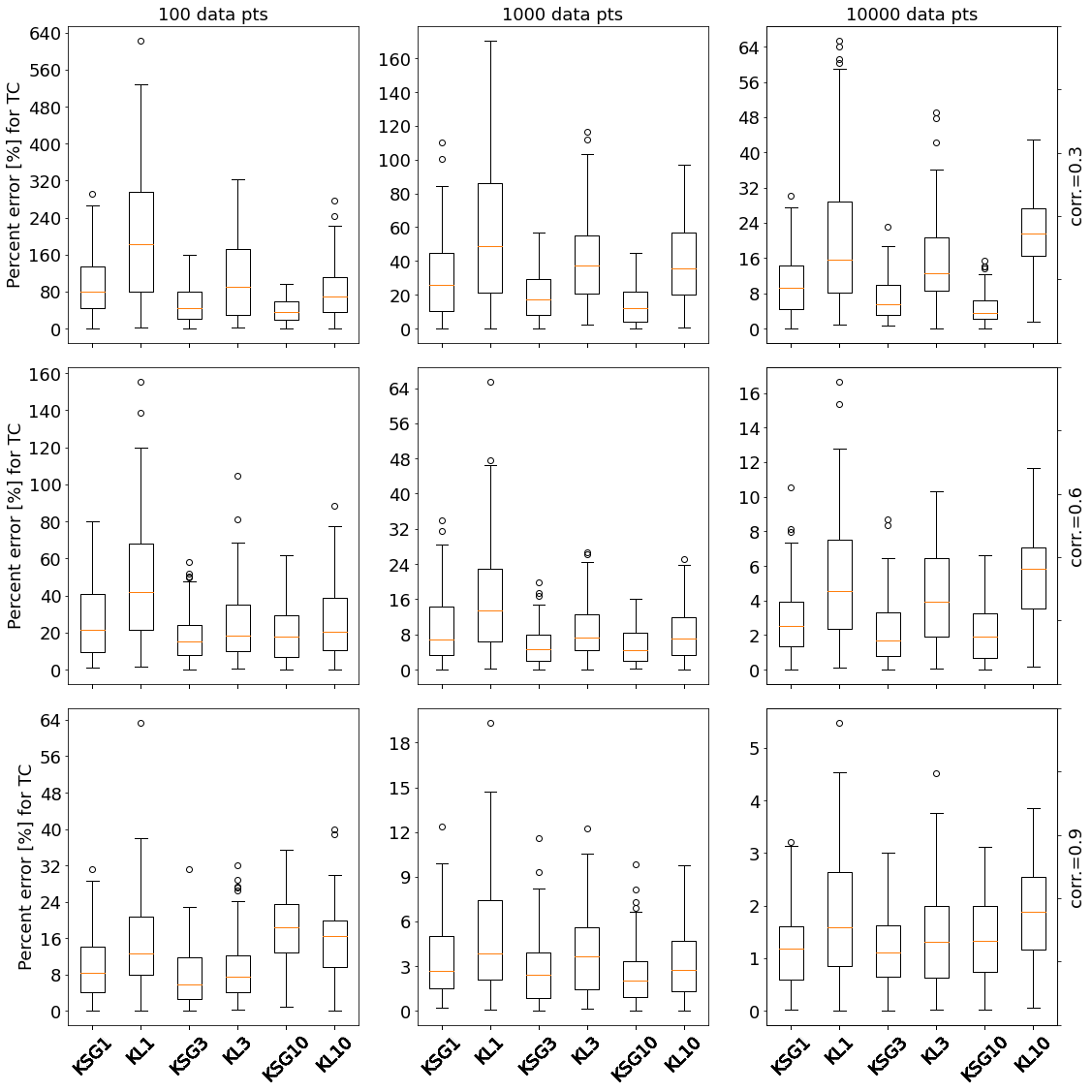

**Figure S7**: Common 3-node network motifs

| **No interaction** | **One edge (Two-genes)** | **Fan-out** | **Fan-in** | **Cascade** | **Feed-Forward-Loop (FFL)** |
| --- | --- | --- | --- | --- | --- |
| 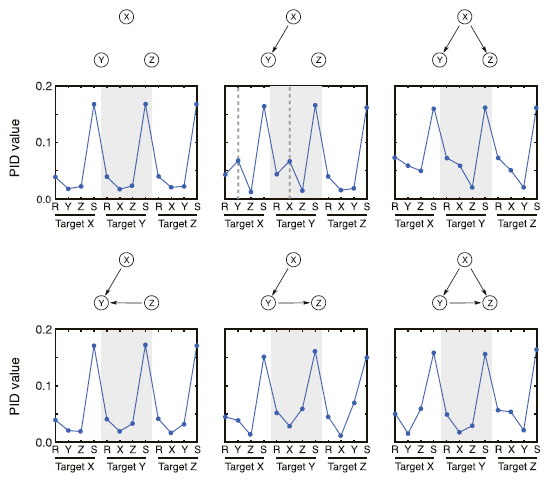 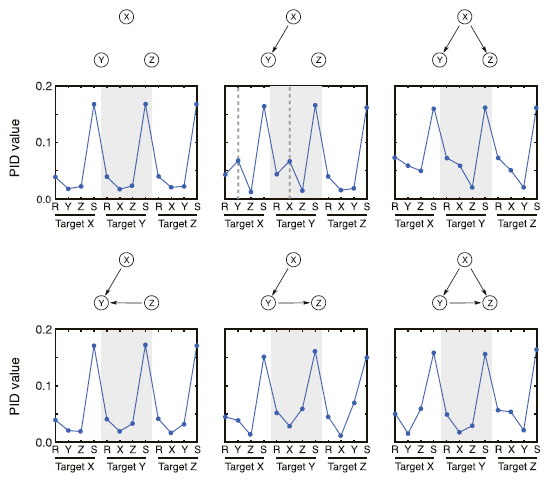 | | | | | |

**Figure S8:** *Sorted boxplots of* percentage AUPR difference (increase or decrease) relative to the gold standard combination [ML,CLR] *for different combinations of MI estimator and GRN inference algorithm for the 6 different Yeast networks from DREAM3. Each boxplot represents 10 replicates.*

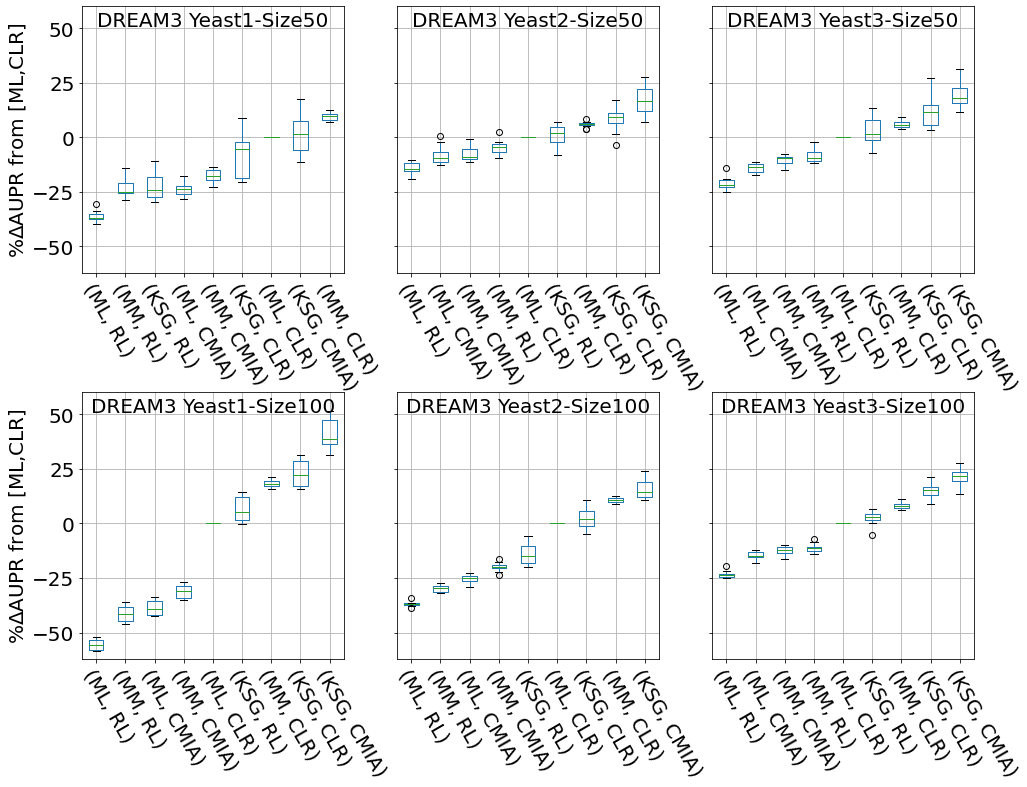

**Figure S9:** *Sorted boxplots of* percentage AUPR difference (increase or decrease) relative to the gold standard combination [ML,CLR] *for different combinations of MI estimator and GRN inference algorithm for the 5 different networks of 100 genes from DREAM4. Each boxplot represents 10 replicates.*

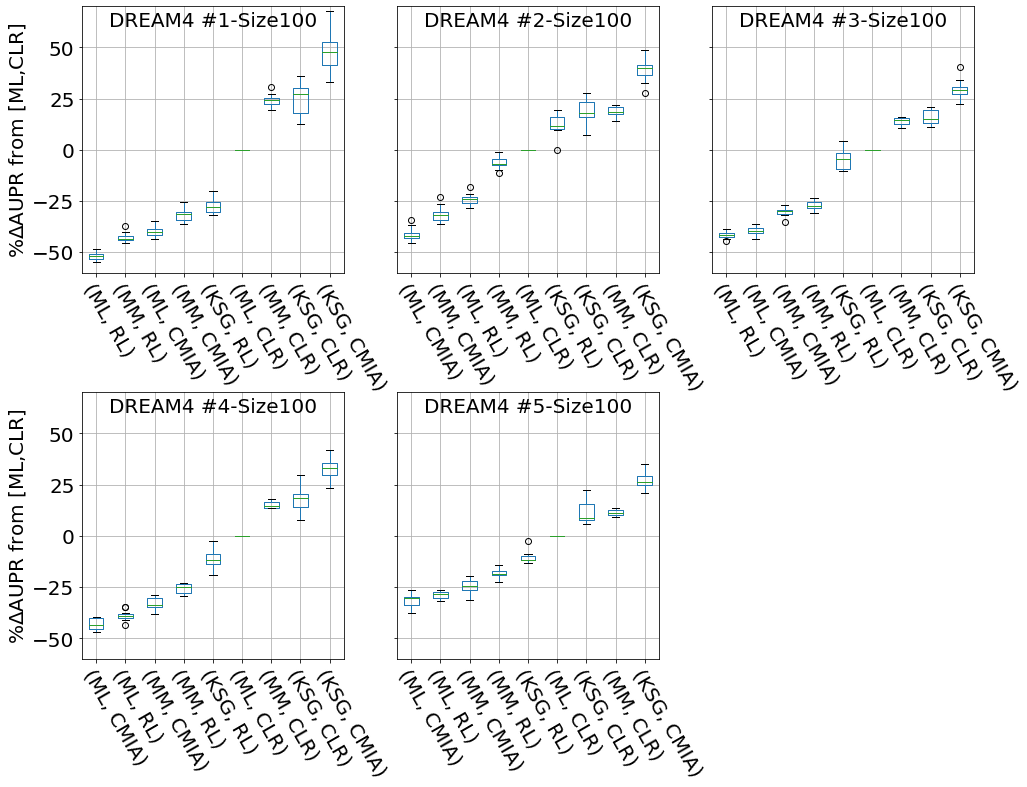

**Figure S10:** Area Under Precision-Recall curve (AUPR) vs. different number of bins or k-neighbors. For the five 50 gene networks from DREAM3, with 10 replicates each, we calculated the AUPR for two inference algorithm and MI estimator {CLR,ML} with blue dots and {CMIA,KSG} with purple dots for different number of bins for ML, and different number of k-neighbors for KSG. The black dashed vertical line represents k=3 and the solid black line represents #bins = floor(sqrt(data_pts)).

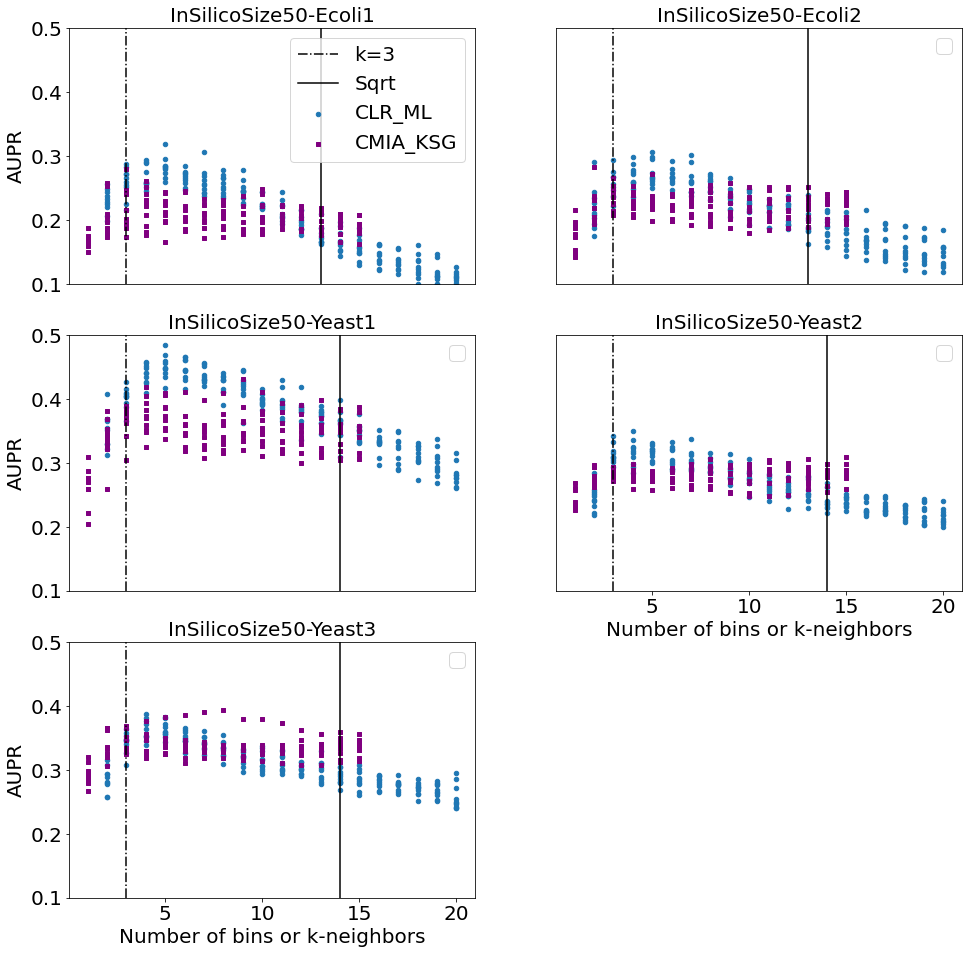

**Figure S11**: The different steps for evaluating GRN inference performance.

Simulating gene expression data

Discretizing/ density estimation

Mutual Information calculations

GRN Inference algorithm

GRN performance evaluation

**Figure S12:** A schematic GRN inference example. The true network contains 10 genes (a.k.a. nodes), and 11 interactions (or edges). The prediction algorithm correctly predicted 6 times (True positive), missed 5 interactions (False negative), and predicted 2 interactions that did not exist (False positive).

**True positive**

**False positive**

**False negative**

G9

G0

G4

G2

G1

G5

G6

G3

G7

G8
