## Additional file 3 for "Gene regulation network inference using k-nearest neighbor-based mutual information estimation-Revisiting an old DREAM"

**Additional file 3: Supplementary information table S1-2**

**Table S1**: Median AUPR values for different combinations of MI estimator and GRN inference algorithm for different network sizes.

| Data set | Network Size | Inf-algo | MI-est | median_AUPR | AUPR_relative |
| --- | --- | --- | --- | --- | --- |
| DREAM3 | 50 | ARACNE | KL | 0.0125 | -95.6782 |
| DREAM3 | 50 | ARACNE | KSG | 0.1355 | -46.5551 |
| DREAM3 | 50 | ARACNE | ML | 0.068 | -72.8548 |
| DREAM3 | 50 | ARACNE | MM | 0.088 | -63.9923 |
| DREAM3 | 50 | CLR | KL | 0.0925 | -64.3731 |
| DREAM3 | 50 | CLR | KSG | 0.2635 | 7.698352 |
| DREAM3 | 50 | CLR | ML | 0.239 | 0 |
| DREAM3 | 50 | CLR | MM | 0.253 | 7.38459 |
| DREAM3 | 50 | CMI2rt | KL | 0.008 | -97.1179 |
| DREAM3 | 50 | CMI2rt | KSG | 0.0675 | -72.6237 |
| DREAM3 | 50 | CMI2rt | ML | 0.01 | -96.1632 |
| DREAM3 | 50 | CMI2rt | MM | 0.013 | -95.0144 |
| DREAM3 | 50 | CMIA | KL | 0.0925 | -62.8254 |
| DREAM3 | 50 | CMIA | KSG | 0.2855 | 16.04475 |
| DREAM3 | 50 | CMIA | ML | 0.2205 | -17.4603 |
| DREAM3 | 50 | CMIA | MM | 0.225 | -10.2612 |
| DREAM3 | 50 | RL | KL | 0.021 | -93.6739 |
| DREAM3 | 50 | RL | KSG | 0.246 | -1.98065 |
| DREAM3 | 50 | RL | ML | 0.206 | -23.5236 |
| DREAM3 | 50 | RL | MM | 0.2315 | -10.7137 |
| DREAM3 | 50 | SA_CLR | KL | 0.084 | -64.9525 |
| DREAM3 | 50 | SA_CLR | KSG | 0.29 | 15.92868 |
| DREAM3 | 50 | SA_CLR | ML | 0.188 | -36.7611 |
| DREAM3 | 50 | SA_CLR | MM | 0.189 | -33.5586 |
| DREAM3 | 100 | ARACNE | KL | 0.018 | -89.3707 |
| DREAM3 | 100 | ARACNE | KSG | 0.103 | -50.4808 |
| DREAM3 | 100 | ARACNE | ML | 0.0515 | -75.2367 |
| DREAM3 | 100 | ARACNE | MM | 0.068 | -64.5277 |
| DREAM3 | 100 | CLR | KL | 0.0625 | -71.6402 |
| DREAM3 | 100 | CLR | KSG | 0.246 | 10.39176 |
| DREAM3 | 100 | CLR | ML | 0.215 | 0 |
| DREAM3 | 100 | CLR | MM | 0.2305 | 17.28043 |
| DREAM3 | 100 | CMI2rt | KL | 0.014 | -93.9775 |
| DREAM3 | 100 | CMI2rt | KSG | 0.0505 | -76.0099 |
| DREAM3 | 100 | CMI2rt | ML | 0.002 | -99.1506 |
| DREAM3 | 100 | CMI2rt | MM | 0.004 | -97.9335 |
| DREAM3 | 100 | CMIA | KL | 0.0735 | -66.5423 |
| DREAM3 | 100 | CMIA | KSG | 0.261 | 22.61162 |
| DREAM3 | 100 | CMIA | ML | 0.142 | -35.2527 |
| DREAM3 | 100 | CMIA | MM | 0.157 | -28.7529 |
| DREAM3 | 100 | RL | KL | 0.0305 | -84.6249 |
| DREAM3 | 100 | RL | KSG | 0.22 | 0.643777 |
| DREAM3 | 100 | RL | ML | 0.105 | -46.7633 |
| DREAM3 | 100 | RL | MM | 0.138 | -29.1302 |
| DREAM3 | 100 | SA_CLR | KL | 0.07 | -66.2411 |
| DREAM3 | 100 | SA_CLR | KSG | 0.2585 | 20.128 |
| DREAM3 | 100 | SA_CLR | ML | 0.073 | -57.0211 |
| DREAM3 | 100 | SA_CLR | MM | 0.077 | -56.5227 |
| DREAM4 | 100 | ARACNE | KL | 0.002 | -99.0589 |
| DREAM4 | 100 | ARACNE | KSG | 0.129 | -43.1122 |
| DREAM4 | 100 | ARACNE | ML | 0.08 | -68.2798 |
| DREAM4 | 100 | ARACNE | MM | 0.103 | -58.4942 |
| DREAM4 | 100 | CLR | KL | 0.03 | -87.9027 |
| DREAM4 | 100 | CLR | KSG | 0.26 | 17.76752 |
| DREAM4 | 100 | CLR | ML | 0.232 | 0 |
| DREAM4 | 100 | CLR | MM | 0.2615 | 15.45322 |
| DREAM4 | 100 | CMI2rt | KL | 0.001 | -99.446 |
| DREAM4 | 100 | CMI2rt | KSG | 0.086 | -64.4488 |
| DREAM4 | 100 | CMI2rt | ML | 0.001 | -99.4778 |
| DREAM4 | 100 | CMI2rt | MM | 0.002 | -99.1038 |
| DREAM4 | 100 | CMIA | KL | 0.031 | -87.329 |
| DREAM4 | 100 | CMIA | KSG | 0.302 | 33.58116 |
| DREAM4 | 100 | CMIA | ML | 0.1445 | -40.0692 |
| DREAM4 | 100 | CMIA | MM | 0.1685 | -30.4858 |
| DREAM4 | 100 | RL | KL | 0.002 | -98.9941 |
| DREAM4 | 100 | RL | KSG | 0.226 | -10.0136 |
| DREAM4 | 100 | RL | ML | 0.159 | -38.7358 |
| DREAM4 | 100 | RL | MM | 0.1905 | -23.9748 |
| DREAM4 | 100 | SA_CLR | KL | 0.031 | -87.3056 |
| DREAM4 | 100 | SA_CLR | KSG | 0.301 | 30.21067 |
| DREAM4 | 100 | SA_CLR | ML | 0.076 | -66.971 |
| DREAM4 | 100 | SA_CLR | MM | 0.082 | -65.1566 |

**Table S2**: Median AUPR values for different combinations of MI estimator and GRN inference algorithm for different organisms.

| Organism | Inf-algo | MI-est | median_AUPR | AUPR_relative |
| --- | --- | --- | --- | --- |
| Ecoli | ARACNE | KL | 0.017 | -88.7796 |
| Ecoli | ARACNE | KSG | 0.0775 | -52.5158 |
| Ecoli | ARACNE | ML | 0.033 | -79.8132 |
| Ecoli | ARACNE | MM | 0.0545 | -66.5278 |
| Ecoli | CLR | KL | 0.043 | -69.9583 |
| Ecoli | CLR | KSG | 0.159 | 5.880966 |
| Ecoli | CLR | ML | 0.1675 | 0 |
| Ecoli | CLR | MM | 0.1965 | 19.87512 |
| Ecoli | CMI2rt | KL | 0.008 | -94.3472 |
| Ecoli | CMI2rt | KSG | 0.0315 | -79.5775 |
| Ecoli | CMI2rt | ML | 0.0025 | -98.5609 |
| Ecoli | CMI2rt | MM | 0.003 | -97.9864 |
| Ecoli | CMIA | KL | 0.0505 | -66.5423 |
| Ecoli | CMIA | KSG | 0.1905 | 19.69697 |
| Ecoli | CMIA | ML | 0.1145 | -33.4421 |
| Ecoli | CMIA | MM | 0.133 | -21.9093 |
| Ecoli | RL | KL | 0.034 | -76.546 |
| Ecoli | RL | KSG | 0.1565 | -2.00302 |
| Ecoli | RL | ML | 0.096 | -38.4287 |
| Ecoli | RL | MM | 0.1245 | -21.7161 |
| Ecoli | SA_CLR | KL | 0.055 | -66.4694 |
| Ecoli | SA_CLR | KSG | 0.1925 | 19.6799 |
| Ecoli | SA_CLR | ML | 0.081 | -53.083 |
| Ecoli | SA_CLR | MM | 0.0865 | -51.4651 |
| Yeast | ARACNE | KL | 0.016 | -94.7018 |
| Yeast | ARACNE | KSG | 0.1545 | -46.5551 |
| Yeast | ARACNE | ML | 0.077 | -72.215 |
| Yeast | ARACNE | MM | 0.1005 | -63.8766 |
| Yeast | CLR | KL | 0.096 | -63.5952 |
| Yeast | CLR | KSG | 0.2915 | 10.4849 |
| Yeast | CLR | ML | 0.2685 | 0 |
| Yeast | CLR | MM | 0.2865 | 8.674749 |
| Yeast | CMI2rt | KL | 0.012 | -95.4876 |
| Yeast | CMI2rt | KSG | 0.0945 | -68.5083 |
| Yeast | CMI2rt | ML | 0.01 | -96.2329 |
| Yeast | CMI2rt | MM | 0.013 | -95.0144 |
| Yeast | CMIA | KL | 0.101 | -63.0312 |
| Yeast | CMIA | KSG | 0.328 | 18.42349 |
| Yeast | CMIA | ML | 0.2235 | -18.074 |
| Yeast | CMIA | MM | 0.236 | -14.5211 |
| Yeast | RL | KL | 0.024 | -92.4062 |
| Yeast | RL | KSG | 0.2515 | 0 |
| Yeast | RL | ML | 0.196 | -27.825 |
| Yeast | RL | MM | 0.219 | -14.0642 |
| Yeast | SA_CLR | KL | 0.0975 | -64.1355 |
| Yeast | SA_CLR | KSG | 0.3225 | 18.01125 |
| Yeast | SA_CLR | ML | 0.1745 | -37.9402 |
| Yeast | SA_CLR | MM | 0.1775 | -36.4665 |

**Table S3**: Characteristics of the 10 synthetic networks from DREAM3 and statistics of the different 3-node network motifs extracted.

| **Network** | **SS data** | **Edges** | **Triplets** | **No Interaction** | **Two-genes** | **Fan-in** | **Cascade** | **Fan-out** | **FFL** | **Sum of 2 edges** | **Sum of 2&3 edges** |
| --- | --- | --- | --- | --- | --- | --- | --- | --- | --- | --- | --- |
| **InSilicoSize100-Ecoli1** | 341 | 125 | 161700 | 150051 | 11059 | 47 | 55 | 477 | 11 | 579 | 590 |
| **InSilicoSize100-Ecoli2** | 322 | 119 | 161700 | 150759 | 10228 | 24 | 51 | 630 | 8 | 705 | 713 |
| **InSilicoSize100-Yeast1** | 401 | 166 | 161700 | 146042 | 15113 | 75 | 212 | 193 | 65 | 480 | 545 |
| **InSilicoSize100-Yeast2** | 401 | 389 | 161700 | 127499 | 30631 | 627 | 1231 | 1361 | 351 | 3219 | 3570 |
| **InSilicoSize100-Yeast3** | 401 | 551 | 161700 | 115759 | 39003 | 1385 | 2052 | 2382 | 1119 | 5819 | 6938 |
| **InSilicoSize50-Ecoli1** | 170 | 62 | 19600 | 16936 | 2361 | 21 | 41 | 232 | 9 | 294 | 303 |
| **InSilicoSize50-Ecoli2** | 169 | 82 | 19600 | 16230 | 2816 | 47 | 20 | 475 | 12 | 542 | 554 |
| **InSilicoSize50-Yeast1** | 201 | 77 | 19600 | 16204 | 3126 | 43 | 103 | 94 | 30 | 240 | 270 |
| **InSilicoSize50-Yeast2** | 201 | 160 | 19600 | 13056 | 5536 | 241 | 306 | 333 | 128 | 880 | 1008 |
| **InSilicoSize50-Yeast3** | 201 | 173 | 19600 | 12629 | 5812 | 195 | 303 | 487 | 174 | 985 | 1159 |
